# Conserved synaptic modifications by humanized tau through diverse isoform- and mutation-specific intracellular interactions

**DOI:** 10.64898/2026.11.03.763730

**Authors:** Emily W. Nickerson, Justice Pomeroy-Tuck, Samia Pratt, Carly Rodriguez, Tengteng Zhao, Tim Stodola, Brian Hoffman, Rick Maser, Erik B. Bloss

## Abstract

Mutations in the microtubule-binding protein tau are considered causal for a diverse group of neurodegenerative diseases collectively referred to as primary tauopathies. Specific tauopathies are associated with different mutations, distinct tau isoforms, manifest in unique cellular and circuit-wide patterns of degeneration, and present with specific clinical symptoms and trajectories. The mechanisms that link variations in the primary tau protein structure to distinct forms of neural circuit dysfunction remain largely unknown. Here, we created an orthogonal viral resource to systematically examine the impact of expressing humanized tau isoforms harboring different disease-relevant mutations in excitatory pyramidal cells from the mouse prelimbic cortex. We used these reagents to show that expression of mutant humanized tau (hTau) as either of two isoforms produced very similar patterns of circuit-specific synaptic disconnection on cortical projection neurons. However, mass spectrometry revealed that the interactomes of these different tau species varied markedly as functions of isoform, mutation, and subcellular location. We validated the functional relevance of increased hTau interactions with mitochondrial proteins in induced neurons differentiated from Ngn2-iPSCs expressing different hTau constructs. These results suggest different disease-relevant hTau proteins likely use distinct mechanisms to drive a highly stereotyped form of synapse loss in cortical neurons.

## Introduction

Primary tauopathies are a diverse collection of neurodegenerative diseases whose clinical progression follows the progressive accumulation of the microtubule binding protein tau^1^. The spatial and cellular pattern of accumulation and molecular identity of the tau proteins found in the insoluble aggregates are distinct among different tauopathies^2^, suggesting a link between primary protein structure, cellular susceptibility, and disease phenotype. Despite this relationship, exactly how different tau primary structures produce dysfunction in cells and neural circuits is poorly understood.

The primary structure of tau depends on the isoform, for which there are six that exist in the human CNS^2^. Each isoform preferentially localizes to different subcellular neuronal compartments^3^, suggesting isoform-specific functional roles within neurons^4^. Disease-relevant mutations within the tau protein also prominently shape intracellular function; for example, the P301L mutation is located within the microtubule-binding domain and dramatically increases tau aggregation^5,6^; A152T, on the other hand, occurs within the proline-rich region and decreases tau aggregation^7^. R406W is in the C-terminus of the protein and produces mixed effects in aggregation assays^5,8^. Although each of these mutations is associated with elevated risk for frontotemporal dementia (FTD)^9-11^, exactly how they modify synaptic connectivity, tau phosphorylation, and tau’s intracellular function remain unclear.

To explore the mechanisms underlying tau’s pathogenetic features and serve as preclinical platforms for therapeutic testing, a variety of transgenic models have been engineered to express wild-type or mutant humanized tau^12-16^. However, the reliance on random transgenic insertion strategies of tau’s coding sequences into the mouse genome can produce artifactual toxicity on its own^17^ and precludes the use of rigorous genetic controls that compare identical expression of wild type vs. mutant tau. Moreover, most models utilize the P301L/S mutation, and no work has systematically examined whether the effects of mutant tau depend on the protein isoform. Last, the focus on neural circuits that are vulnerable specifically in Alzheimer’s disease, which is confounded by the presence of a second neuropathology, has limited mechanistic insights of the development and progression of primary tauopathies.

Here, we built a viral toolkit to examine the impact of expressing the 0N4R and 2N4R isoforms of human tau (hTau), either as wild-type (WT) or containing the disease-linked A152T, P301L, or R406W mutations. Our viral approach enabled hTau expression to be spatially and temporally restricted to subsets of excitatory projection neurons in the mouse prelimbic cortex, a region considered functionally and anatomically analogous to human frontal cortical regions that are vulnerable in FTD^18-20^. Each of the disease-linked hTau isoform-mutation pairings markedly decreased excitatory synaptic connectivity and altered synapse structural morphology. Although these changes suggest uniform neuronal responses, subcellular mass spectrometry analysis revealed dramatic differences between each hTau’s protein-protein interactions. A152T and P301L’s interactomes revealed little overlap between isoform and subcellular compartments: at somatodendritic sites, these variants primarily interacted with extracellular matrix proteins while at long-range axonal projection sites engaged with proteins important for the regulation of gene expression. Conversely, R406W’s interactome exhibited similar interactions with mitochondrial proteins across both subcellular compartments. Using hTau-expressing Ngn2-differentiated neurons made from human induced pluripotent stem cells, we found both hTau^WT^ and hTau^R406W^ altered the transcriptional regulation of mitochondrial-associated genes and impacted mitochondrial capacity.

Overall, these results suggest that different disease-linked hTau mutations uniformly produce synaptic disconnection and remodeling but through disparate mechanisms shaped by the predominant isoform expressed in each cellular compartment. Collectively, these dendritic and –omics datasets are a resource that can be used by the scientific community to generate new hypotheses about isoform-mutation relationships and the engagement of each tau species with specific elements of disease-relevant pathways. Moreover, the reagents described here can be easily modified to include other hTau mutations and isoforms and can be used with cell-type or circuit-level specificity to test these hypotheses in a rigorous and controlled manner.

## RESULTS

### Viral strategy to drive cellular and circuit specific expression of human tau

To restrict the expression of the hTau protein to a single circuit and cell type, we made several modifications to existing hTau viral constructs^21^. We inverted the coding sequence and added loxP sites to render the construct Cre-dependent, utilized a human Synapsin promoter to promote stable long-term neuronal expression, and packaged these in AAV2/1 (**Figure 1A-B; Figure S1A)**. Each hTau construct expressed either the 0N4R or 2N4R isoform of WT or one of three disease-related mutations each located in a different functional domain of the hTau protein: A152T, P301L, or R406W. Last, we added a linker sequence and N-terminal FLAG tag to the hTau protein, permitting easy visualization and isolation of expressed hTau protein. By pairing the hTau constructs with AAV-mediated CaMKII-Cre expression, we restricted expression of constructs to excitatory neurons in the mouse prelimbic cortex (PL; **Figure 1C**), an understudied region in tauopathy research despite the susceptibility of analogous human brain to tau accumulation in FTD and other primary tauopathies^22,23^. Postembedding immunogold labeling identified FLAG-labeled gold particles at perisomatic locations associated with microtubules and in postsynaptic locations within dendritic spines (**Figure 1C**), similar to what has been previously reported^16^. After three months of expression, there was a strong correlation between EGFP intensity and hyperphosphorylated hTau (pSer202/pThr205, detected with AT8^+^) in PL neurons across all constructs (0N isoform, **Figure 1D**). Similarly, expression of each construct in PL produced local activation of the microglial network as evidenced by the presence of CLEC7a^+^ cells, a signature of disease-associated microglia^24^ (**Figure S1B-S1C**).

**Figure 1.**
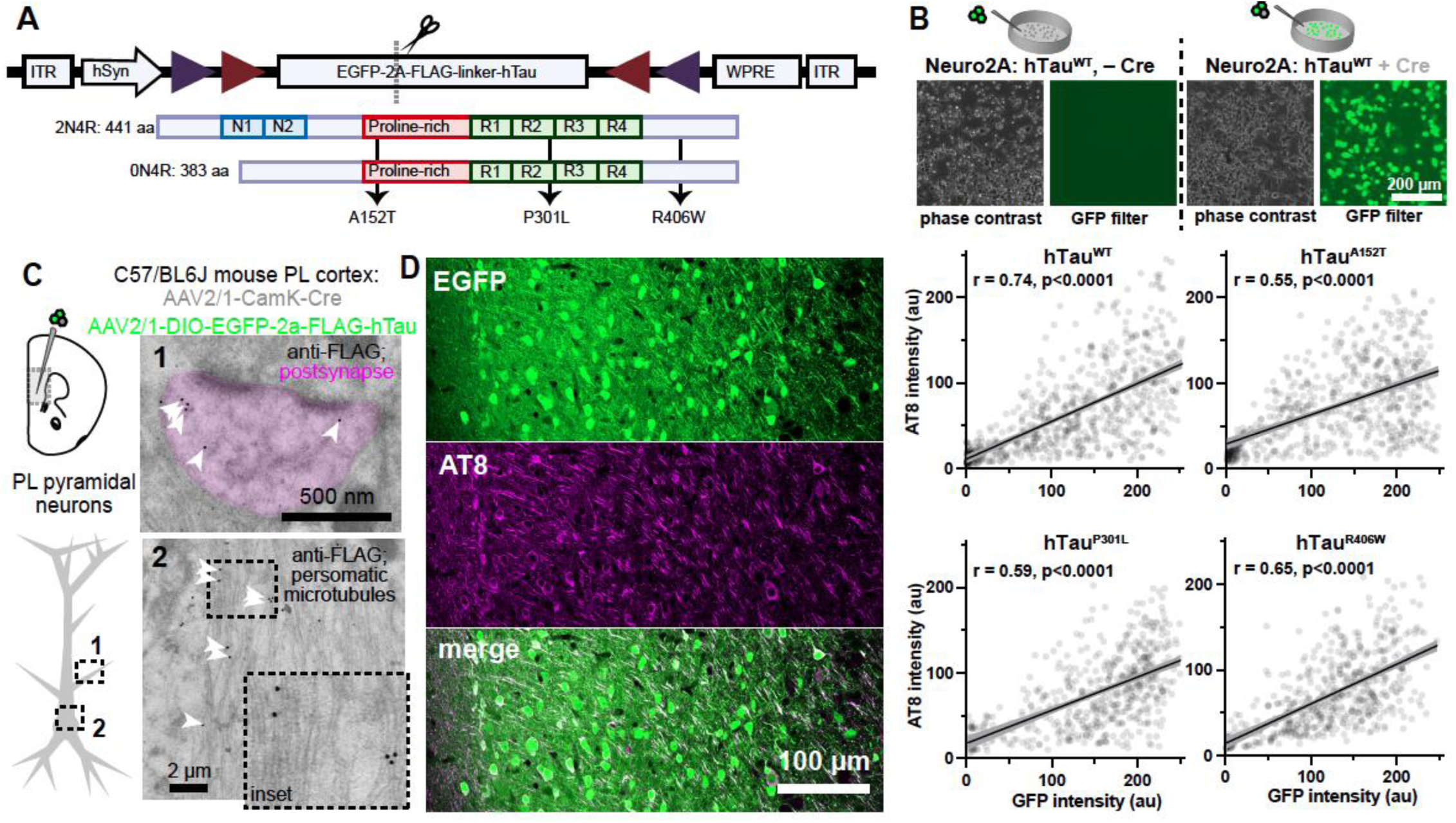
Viral approach to express hTau in mouse prefrontal cortical neurons. **A)** Construct design to drive stable viral Cre-dependent neuronal expression of EGFP-2a-FLAG-hTau as 0N4R or 2N4R isoforms and containing one of four sequences: hTau^WT^, hTau^A152T^, hTau^P301L^, and hTau^R406W^. **B)** Phase contrast and EGFP-2a-hTau signals from Neuro2A cells in the absence (left) or presence (right) of viral Cre recombinase. **C)** Postembedding immunogold electron microscopy of FLAG-hTau (white arrowheads) revealed synaptic (pink compartment) and microtubule-associated labeling of virally expressed hTau (0N4R hTau^P301L^) within prelimbic excitatory cortical neurons. D) Correlated expression of EGFP-2a-hTau and hyperphosphorylated hTau (AT8; pSer202/205) in prelimbic cortex across 0N4R hTau^WT^, 0N4R hTau^A152T^, 0N4R hTau^P301L^, and 0N4R hTau^R406W^.

### Compartment-specific decreases in synaptic density by mutant hTau

Dendritic spine loss is tightly correlated to cognitive and behavioral decline in tauopathies^25,26^ though there is less definitive data from tauopathies associated with hTau mutations^27-31^. In transgenic mouse models harboring the P301S mutation, such loss is thought to precede insoluble tau aggregation and is considered a key early step towards information processing impairments in neural circuits^32,33^. However, the global expression of mutant hTau in such models precludes identifying whether such synapse loss is driven by pre- or postsynaptic neuronal expression and it remains unknown if synaptic alterations present in P301S transgenic mice generalize to different hTau isoforms and mutations. We leveraged our viral constructs to restrict the expression of hTau to layer 2/3 PL neurons projecting to the basolateral (PL-to-BLA) by pairing bilateral injections of AAVretro-Cre in the basolateral amygdala with bilateral PL injections of an AAV expressing an hTau construct or a control fluorophore (EGFP; **Figure 2A; Table S1**). Such an approach leverages a within-animal control while also allowing the same excitatory projection neurons to be sampled across mice.

**Figure 2.**
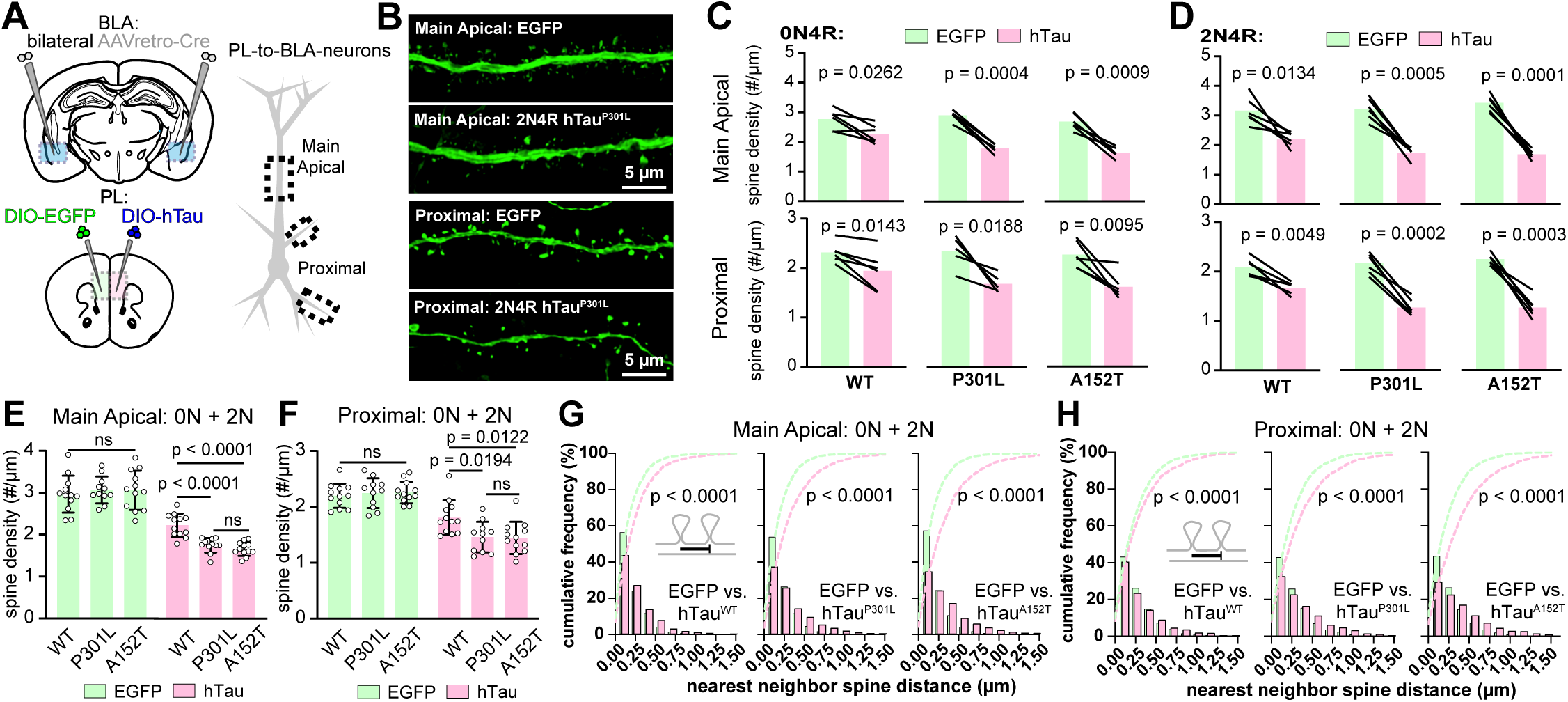
Mutant hTau drives dendritic spine loss on PL-to-BLA neurons. **A**) Dual viral injection strategy to drive EGFP (control) or EGFP-2a-hTau expression in different hemispheres within the same mouse. **B)** Example segments from EGFP or EGFP-2a-hTau-expressing main apical dendrites (top) or proximal apical oblique or basal dendrites. **C)** Within-animal dendritic spine densities between branches from EGFP (control) and 0N4R hTau constructs, with data from main apical branches (top) and proximal apical oblique or basal branches (bottom). D) same as in (**C**) but for 2N4R constructs. **E**) Across-animal comparisons for EGFP and EGFP-2a-hTau-expressing main apical branches. **F**) Same as in (**E**) but for proximal branches. **G)** Cumulative frequency plot and bar plot of nearest neighbor spines along main apical branches between spines from EGFP and EGFP-2a-hTau branches (0N and 2N isoforms combined; see Figure S2 for each isoform). **H)** Same as in (**G**), but for proximal apical oblique or basal dendrites (see Figure S2 for each isoform).

The dendritic compartments of cortical pyramidal cells have significantly different electrical and structural properties that shape the integration of synaptic input^34^. To determine how the different hTau proteins modified synaptic connectivity across these locations, we used confocal microscopy to reconstruct dendritic spine densities across the distal apical tuft, the main apical trunk, and proximal apical oblique/basal branches (**Figure 2B**). Immunolabeling of the FLAG tag showed that the viral expression of each construct was restricted to the targeted hemisphere (**Figure S2A**). Compared to the spine densities from the EGFP hemisphere of each mouse, the expression of each hTau construct (both the 0N4R and 2N4R isoforms, and including hTau^WT^), significantly reduced spine densities on main apical and proximal dendrites (**Figure 2C-2D; Figure S2A-S2C**). As expression of EGFP produced replicable dendritic spine densities across mice from each group (**Figure 2E-2F**), we also performed across-animal comparisons to examine the relative effect size across hTau mutations. Spine densities were significantly lower on the main apical and proximal branches from hTau^P301L^- and hTau^A152T^-expressing neurons (∼40% decrease along the main apical trunk and ∼33% on the proximal branches) relative to those expressing to hTau^WT^ (∼25% decrease along trunk and 20% on proximal branches; **Figure 2E-2F**). In contrast, the spine densities on the distal tuft dendrites were only marginally affected by hTau (**Figure S2D**), suggesting circuit-specific synaptic responses to hTau mutations.

The spatial clustering of spines is a strong determinant of their ability to cooperatively depolarize the parent dendrite^35^ and influence across-synapse plasticity^36^, so we examined the spatial distribution of the remaining spines along our reconstructed branches. Using a nearest neighbor analysis, we found that each hTau construct led to a significant increase in the distance between dendritic spines on the main apical and proximal dendritic branches (**Figure 2G-2H**), suggesting that spine loss was distributed along each dendritic segment rather than localized to discrete sites along the branch. While the increase of nearest neighbor distances was evident with hTau^WT^, the increase in distances suggested spines on hTau mutant-expressing neurons were nearly twice as far apart from each other on the main branch and 50% further apart on proximal branches. Consistent with the weaker effect on spine density, this effect was much weaker in the distal tuft dendrites (**Figure S2E-S2F**). The decreased vulnerability of hTau-mediated disconnection of distal tuft dendritic synapses could reflect the receipt of inputs from different afferent circuits or the distinct synaptic strategy used on these branches (**Figure S2G**). Collectively, these results show that both the 0N4R and 2N4R isoforms of hTau^P301L^ and hTau^A152T^ significantly decrease synaptic densities and local dendritic clustering selectively along apical and proximal dendrites of prelimbic cortical neurons.

### Mutation- and isoform-specific modifications of dendritic spine morphology

Dendritic spine structure is tightly correlated with synapse function^37^. Two morphological spine features – spine neck length and spine head volume – strongly influence synaptic electrical compartmentalization^38^ and synaptic weight^39^, respectively. Exactly how different hTau proteins impact synapse structure is poorly understood, so we reconstructed the spine neck lengths and spine head volumes of individual spines (**Figure 3A-3B**). Along the main apical dendrite, dendritic spines from hTau expressing cells had significantly shorter neck lengths than EGFP-expressing synapses on branches from the same locations (**Figure 3C-3**). These effects were significantly greater for hTau^P301L^ and hTau^A152T^ mutations compared to hTau^WT^ and were identical across the 0N and 2N isoforms (**Figure S3A-B**). Effectively identical results were found on the proximal apical/basal branches **(Figure S3C-E)**. Like spine density, these effects were much weaker on spines from tuft branches (**Figure S3F-H**). In contrast, the effect of hTau expression on dendritic spine head volume was much weaker: only hTau^A152T^ produced a significant decrease in spine head volume (**Figure 3F-3H**), which was restricted to the main apical branch **(Figure S3I-S3N)**.

**Figure 3.**
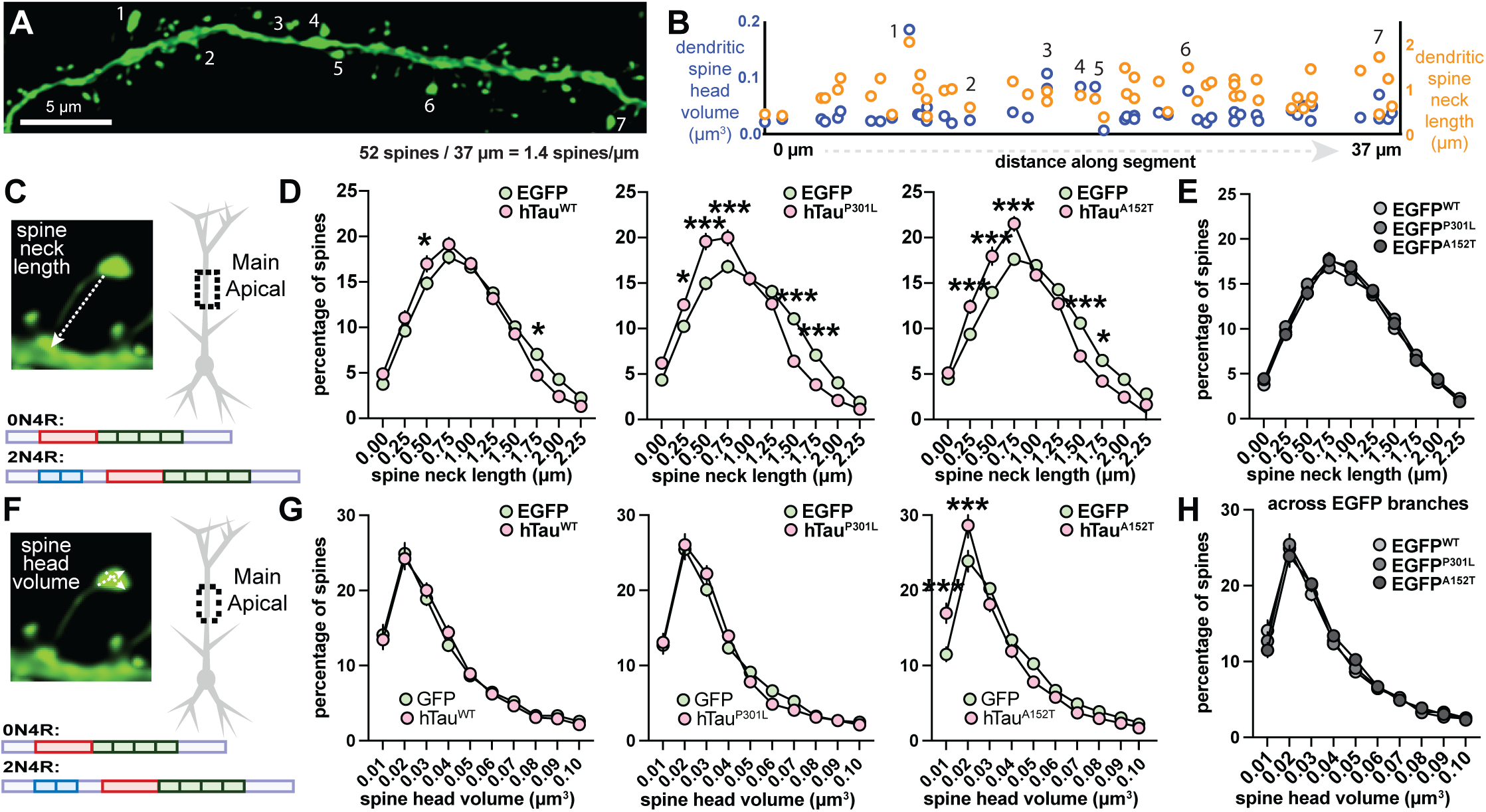
Disease-linked hTau mutations predominantly alter spine neck morphology. **A)** Spine morphologies vary significantly along a dendritic segment. **B**) Individual neck lengths and head volumes from spines reconstructed along the segment. **C**) Spine neck lengths, measured from the center of the spine head to the closest node of the dendritic cable, were taken from segments along the main apical branch from EGFP (control) or EGFP-2a-hTau constructs (30 branches from 0N4R and 30 branches from 2N4R; combined) from PL-to-BLA neurons. **D**) Spine neck length distributions were shifted towards smaller lengths for hTau each group between EGFP and hTau^WT^, EGFP and hTau^P301L^, and EGFP and hTau^A152T^ (plotted from right to left; 0N and 2N isoforms combined; see Figure S3 for each isoform). **E**) Spine neck length distributions were unchanged between EGFP control mice across these groups. **F-H**) Same as in (**C-E**) but for spine head volume.

Collectively, these results suggest that the most prominent effect of hTau on cortical neuronal spine morphology is on neck length, which was reduced almost uniformly across hTau mutants. From a spine dynamics perspective, the decrease in spine densities and reduction in neck lengths are consistent with increased rates of spine loss relative to spine formation^40^, suggesting that hTau mutations alter spine formation and stability in vivo. As the spine neck controls spine voltage changes and decreases in spine neck length are presumed to correspond with decreases in the neck resistance^41,42^, the changes we see here are likely associated with substantial reductions in spine head excitatory postsynaptic potentials. Whether such a reduction is sufficient to limit the activation of voltage-gated Na^+^, Ca^2+^ ^43,44^ could be addressed in future work.

Last, we asked whether expression of these constructs modified PL neuronal excitability. Past work has suggested that mutant hTau^P301L^ expression reduces neuronal excitability in CA1 neurons^45^ and in entorhinal and parietal cortical neurons^46,47^ while other work has shown hTau^A152T^ increases excitability^12,48^. We drove EGFP or the 0N4R forms of hTau^WT^, hTau^P301L^, hTau^A152T^, or hTau^R406W^ in excitatory PL neurons by pairing Cre-dependent constructs with local CaMKII-Cre injections and we made whole-cell patch clamp recordings from EGFP^+^ superficial neurons in acute slices after 3 months of viral expression (n=74 cells from 26 mice; See Methods). Somatic step current injections were used to determine spiking behavior and ramp current injections were used to identify rheobase. While each hTau construct reduced spike output at 100 pA and 150 pA somatic current injections relative to EGFP-expressing neurons, rheobase did not change across the groups **(Figure S3O-S3P)**. Collectively, these results show that mutant hTau expression drives robust yet selective excitatory synaptic loss, a consistent form of synaptic morphological remodeling, and a reduction of somatic spike output.

### Subcellular location and isoform control hTau protein-protein interactions

Our synaptic reconstructions suggest a convergent mechanism by which distinct disease-relevant hTau mutations drive almost identical forms of synaptic disconnection on PL neurons. Such a mechanism could be implemented by different hTau mutants interacting with similar sets of target proteins. To explore this issue, we paired our viral strategy with tandem mass spectrometry (**Figure 4A**). We drove expression of our hTau constructs in excitatory PL neurons by co-injecting AAV2/1-CaMKII-Cre and we isolated hTau-bound proteins by co-immunoprecipitation of the FLAG-tagged hTau protein and its binding partners. Because we could directly obtain hTau binding proteins from within the heterogenous tissue, we separately microdissected the somatodendritic compartment (located in PL) and the long-range axonal compartment (located in BLA) to gain insight into whether mutant hTau had subcellular specific-interactomes. We multiplexed our samples with tandem mass tags and compared the relative abundances to a control group consisting of mock samples from non-transduced animals and IgG control samples that utilized a non-specific IgG antibody for the immunoprecipitation (**Table S2**). From these same samples, we enriched for phosphopeptides and examined changes in the phosphorylation of hTau (**Figure S4**), which is an established molecular marker of disease progression in tauopathies^49-51^.

**Figure 4.**
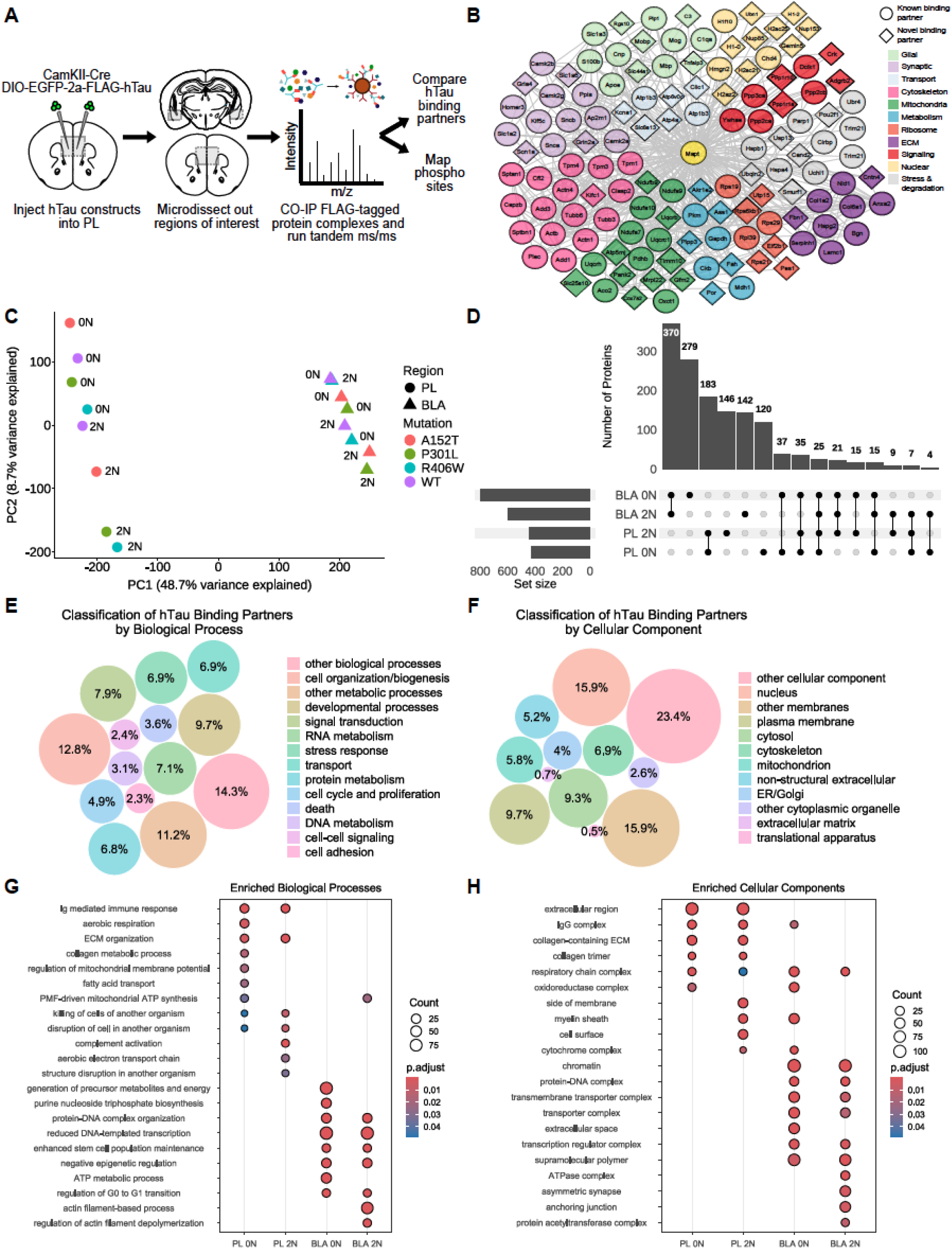
hTau’s interactome is strongly influenced by its intracellular location and primary structure. **A)** PL and BLA were isolated from the brains of mice injected with different AAV hTau constructs. FLAG-tagged hTau and its binding partners were co-immunoprecipitated and analyzed with tandem ms/ms. **B)** Network map, created with String-db and Cytoscape, highlights a selection of the 1,408 proteins that were significantly bound by at least one hTau protein. The proteins belong to a variety of biological processes and cellular components. **C)** Principal component analysis (PCA) of the hTau samples based on grouped abundances. The samples cluster foremost by anatomical region (PC1: 48.7%). The second principal component only accounts for 8.7% of the variance, although the 0N and 2N isoforms do show clear separation in PL. **D)** Upset plot highlights the minimal overlap in binding partners of the 0N4R and 2N4R isoforms in the somatically enriched PL vs axonally enriched BLA. A comparable number of proteins were identified in the different samples with 449 proteins unique to PL (31.9%), 791 unique to BLA (56.2%), and 168 proteins shared amongst the two regions (11.9%). **E)** Proteins were classified according to the MGI GO Slim Categories. Circular packing graph indicates the top biological processes that hTau’s binding partners belonged to. *Other biological process* encompasses anything that doesn’t fit into the other categories. **F)** Circular packing graph representing the top cellular components that hTau’s binding partners belonged to. Neuron-specific structures fall within the *other cellular component* category. **G-H)** Dot plot depicting the significantly upregulated biological processes (**G**) or significantly upregulated cellular components (**H**) for the binding partners of 0N4R or 2N4R hTau proteins compared to control in PL or BLA. The size of the dots represents the number of proteins associated with the pathway and the color scale indicates the adjusted p-value.

Across both cellular compartments, we identified 1,408 total proteins that interacted with hTau, which included both known and novel binding partners (**Figure 4B**) taken from a reference dataset consisting of 14 previously published studies that incorporate a range of different models and methodologies^4,49,52-63^ (**Table S3**). 425 proteins (∼30%) were known binding partners of hTau with 279 additional proteins (∼20%) sharing a close homology to an identified interactor. Among the known identifiers, were many hTau canonical binding partners, including those important for the organization of the cytoskeleton (Dpysl3, Tubb3, Actn1, Add1, Clasp2, Dclk1, Actb, etc.), synaptic plasticity (Camk2a, Ppp3ca, Sptan1, Dlg4, Snca, etc.), cellular metabolism (Ckb, Pkm, Pdhb, Acat1, Aco2, etc.), and regulating tau’s folding and post-translational states (Ppp2ca, Ppp2cb, Ppp5c, Hspa4, Ppia, etc.) (**Table S3**). 704 proteins, or approximately half of the dataset, were novel interactors with hTau. Of the total 1,408 identified proteins, only 168 (12%) were found in both PL and BLA samples, revealing a strong division between hTau interactions across these two compartments (**Figure 4C-4D**). The presence or absence of the two N-terminal repeats also affected hTau’s interactome but to a lesser degree as about half of the total proteins (675 of 1408) were shared across both isoforms. The remaining proteins were split among 0N4R only (31%; 436 of 1408) and 2N4R only (21%; 297 of 1408).

The identified hTau binding proteins belonged to a wide range of biological processes (**Figure 4E**) and localized to several cellular compartments (**Figure 4F**). Using an overrepresentation analysis, we identified the top biological processes (**Figure 4G**) and cellular components (**Figure 4H**) that were significantly upregulated in the 0N4R or 2N4R hTau samples compared to control samples. Within PL, there was a marked convergence in the interactomes of the 0N4R and 2N4R proteins (**Table S4**), such that binding partners associated with the immune system, aerobic respiration, and the extracellular matrix (ECM) were overrepresented. Proteins associated with mitochondrial function and those associated with the ECM were also found in hTau interactomes from the BLA, but to a lesser degree. A similar trend was seen within BLA: both 0N4R and 2N4R hTau bound proteins important for gene regulation and cell cycle phase transitions. Yet there were also notable differences between the interactomes of the two isoforms: the enrichment in energy metabolism was unique to the 0N4R hTau proteins while conversely the 2N4R hTau proteins had increased interactions with actin-binding proteins. Together, these results suggest that both isoform and subcellular locations strongly control how hTau proteins engage with cellular processes.

### The specificity of hTau’s interactome within PL

We next examined the specificity of hTau protein-protein interactions within PL. Out of the 617 total proteins identified, 347 (56.2%) of them were found to be enriched in the interactomes of at least two different hTau proteins, suggesting a high degree of shared functionality across the distinct isoforms and mutations. 176 proteins (28.5%) were bound exclusively by 0N4R hTau proteins, 191 proteins were bound exclusively by 2N4R hTau proteins (31%), and the remainder shared among the isoforms (250 of 617; 40.5%) (**Figure S5A; Table S5**). Overall, the proteins that bound hTau in PL were largely involved with extracellular matrix remodeling, cell killing processes, activation of an immune response, and aerobic respiration (**Figure 5A**; **Figure S5B**). These patterns are consistent with past work identifying functional roles for tau in metabolism and the ECM^59,64,65^. Given hTau’s expression significantly altered dendritic spine composition and these samples are enriched from the somatodendritic compartment, there was a surprising lack of proteins typically associated with synaptic remodeling.

**Figure 5.**
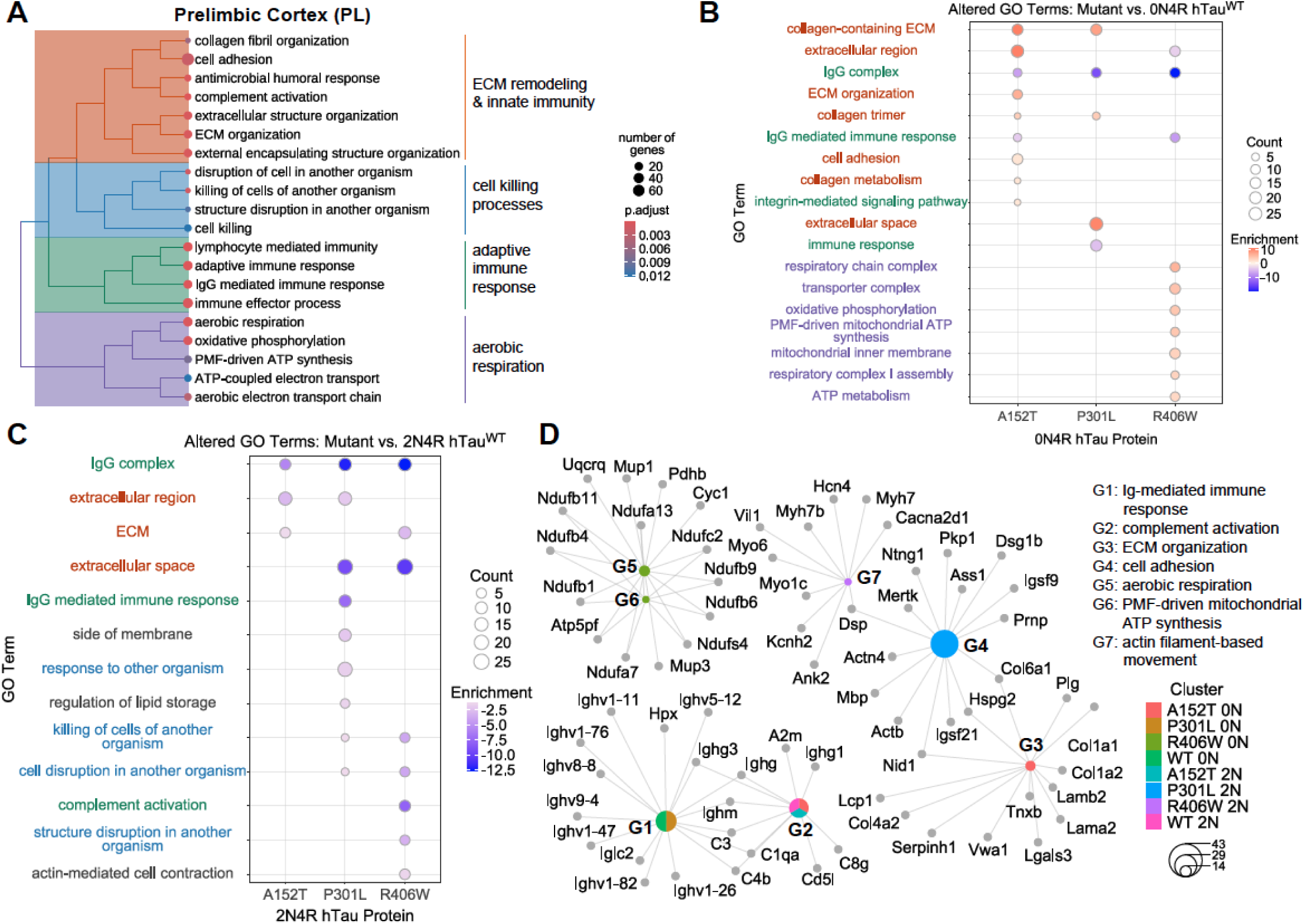
hTau proteins exhibit distinct interactomes in PL. **A)** Hierarchical clustering of key biological processes enriched for in the binding partners of hTau proteins in PL vs control. The size of the nodes denotes the number of genes associated with the GO term, while the color scale indicates the adjusted p value. The GO terms in the treeplot form four major clusters. **B)** Dot plot indicates the top upregulated GO terms relating to biological process and cellular components for the mutant 0N4R hTau proteins compared to 0N4R hTau^WT^. The size of the dots represents the number of proteins associated with the pathway and the color scale indicates the adjusted p-value. **C**) Dot plot indicates the top altered GO terms relating to biological processes and cellular components for the mutant 2N4R hTau proteins compared to 2N4R hTau^WT^. GO terms that were significantly enriched for in the mutant protein’s interactomes are depicted in red, while those significantly downregulated compared to hTau^WT^ are shown in blue. **D)** Gene-concept network constructed from the over-representation analysis comparing the interactomes of the hTau protein to control. The top GO terms (G1-G7) relating to biological processes and the proteins enriched in these pathways are shown. The size of the circle indicates the total number of proteins associated with the GO term.

When comparing the binding partners of the different mutant hTau proteins, a clear divide emerged between mutation and isoform that was evident from the upregulated GO terms of mutant 0N4R proteins (**Figure 5B**). While the interactome of 0N4R hTau^R406W^ was overwhelmingly dominated by mitochondrial proteins and those associated with energy and metabolism, 0N4R hTau^A152T^ and 0N4R hTau^P301L^, bound a significant number of proteins involved in the organization of the ECM. The mitochondrial proteins found within 0N4R hTau^R406W^ spanned multiple different electron transport chain complexes and included Timm9 and Timm10, two key chaperones in the mitochondrial intermembrane space. The variety and type of mitochondrial proteins present, many of which localize to the inner mitochondrial membrane and matrix, strongly suggest that these are not transient interactions with outer mitochondrial proteins. The diversity of ECM proteins detected, and strength of enrichment exhibited by 0N4R hTau^A152T^, and to a lesser extent 0N4R hTau^P301L^, strongly suggests that these mutant proteins were secreted from the cell and accumulated within the extracellular space.

In contrast to the 0N4R results, there were almost no significantly upregulated processes associated with mutant 2N4R hTau proteins compared to 2N4R hTau^WT^ from PL. The significantly downregulated GO terms included pathways relating to the ECM and immune system (**Figure 5C**). However, the mutant 2N4R hTau proteins did have significantly enriched GO terms associated with them relative to control samples (**Figure 5D**), and all four bound a significant number of proteins associated with the ECM. Among the 2N4R mutations, hTau^P301L^’s interactome was enriched with proteins relating to aerobic respiration and the myelin sheath, while hTau^R406W’^s interactome revealed increased proteins relevant to actin dynamics. Collectively, our results from PL reveal isoform- and mutation-specific binding patterns that indicate a surprisingly high degree of functional specialization embedded in hTau interactomes.

### Interactomes of hTau mutants at long-range axonal sites

Given the diversity of hTau isoform and mutant protein-protein interactions within PL, we next examined hTau protein-protein interactions within the BLA, a major axonal target of long-range connections from PL. As in PL, there was strong overlap among the binding partners of the hTau proteins in the BLA, with 64% of the total proteins detected being enriched in the interactomes of at least two hTau species (613 of 959; **Table S6**). The proportion of proteins unique to 0N4R (38.1%; 366 of 959), unique to 2N4R (16.9%; 162 of 959), and shared among the isoforms (45%; 431 of 959) was also comparable to PL.

As expected, proteins critical for cytoskeletal organization and synaptic function were associated with both 0N4R and 2N4R hTau (**Figure S6A**; **Table S6**). For both isoforms, the interactions with actin associated proteins was greater than that of tubulin, consistent with the notion that hTau binding to f-actin^66^ might shape function at presynaptic, actin-rich sites. There were three major processes that were significantly upregulated in the interactomes of the hTau proteins in the BLA (**Figure 6A; Figure S6B**), including a surprisingly robust enrichment for nuclear proteins, especially those involved in DNA accessibility, DNA repair, and gene regulation. Additionally, there was an enrichment in proteins essential for energy generation and metabolism, similar to what was observed in PL, as well as proteins involved in regulating the cell cycle. Proteins linked to cell cycle-related GO terms were overwhelmingly associated with the ATP-dependent BRG1/BRM associated factor chromatin remodeling complex (e.g., Smarca4, Smarca2, Smarcb1, Smarcc1, Smarce1, Arid1a, among many others), which has been shown to play a role in synaptic plasticity and long-term memory formation^67^. Although we were unable to visualize FLAG^+^ proteins within BLA nuclei, the presence of classical nuclear proteins associated with gene regulation and cell cycle control strongly suggest mutant hTau undergoes transcellular transport out of the axon and into recipient cells (similar to^68^).

**Figure 6.**
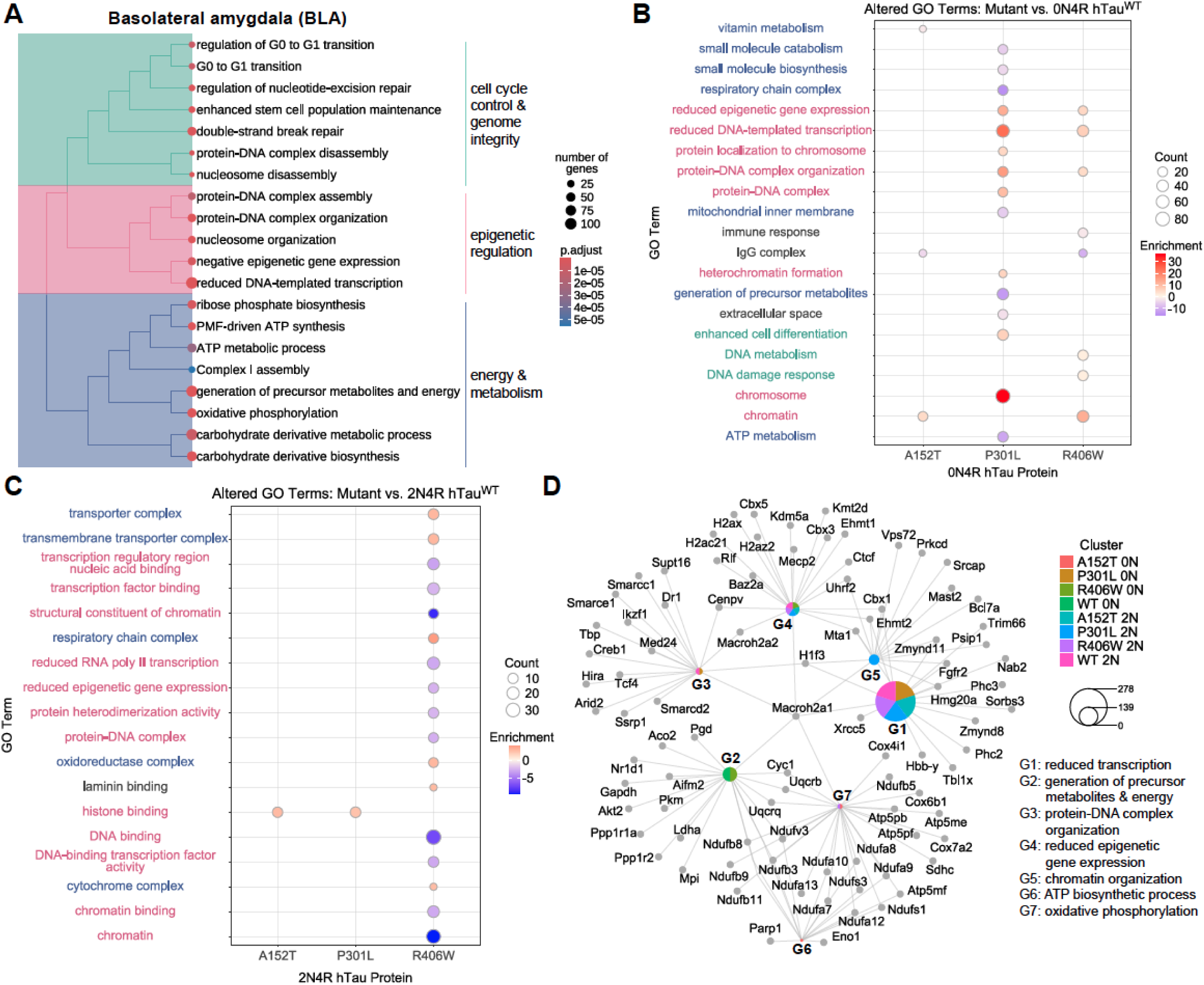
BLA interactome is enriched in proteins that are important for genomic regulation. **A)** Hierarchical clustering of key biological processes enriched for in the binding partners of hTau proteins in BLA vs control. The size of the nodes denotes the number of proteins associated with the GO term, while the color scale indicates the adjusted p value. The GO terms in the treeplot form three major clusters. **B-C)** Dot plot indicates the top upregulated (red) and downregulated (blue) GO terms for the mutant 0N4R hTau proteins compared to 0N4R hTau^WT^ (**B**) or for the mutant 2N4R hTau proteins compared to 2N4R hTau^WT^ (**C**). The size of the dots represents the number of proteins associated with the pathway and the color scale indicates the adjusted p-value. **D**) Gene-concept network constructed from the over-representation analysis comparing the interactomes of the hTau protein to control. The top GO terms (G1-G7) relating to biological processes and the proteins enriched in these pathways are shown. The size of the circle indicates the total number of proteins associated with the GO term.

As in PL, hTau protein-protein interactions in BLA revealed distinct differences between the 0N4R and 2N4R isoforms. 0N4R hTau^P301L^ and hTau^R406W^ interacted with a surprising number of histones and proteins associated with DNA damage response and chromatin remodeling (**Figure 6B**), and the upregulated GO terms suggested a strong and specific pattern of gene repression (GO terms include “reduced epigenetic gene expression”, “reduced DNA-templated transcription”, and “heterochromatin formation”; **Table S6**). 0N4R hTau^P301L^ also exhibited a marked decrease of proteins associated with different metabolic processes compared to 0N4R hTau^WT^. 0N4R hTau^A152T^ exhibited an interactome similar to 0N4R hTau^WT^, both interacting with a substantial number of mitochondrial proteins. In the 2N4R form, hTau^R406W^ had a profile very similar to what was seen with its 0N4R counterpart in PL (**Figure 6C**) with significant upregulation in GO terms associated with aerobic respiration and cellular metabolism. 2N4R hTau^R406W^ lacked interactions with proteins associated with gene expression, contrasting directly with the interactions identified for 0N4R hTau^R406W^. The interactomes of 2N4R hTau^P301L^ and 2N4R hTau^A152T^ largely resembled 2N4R hTau^WT^ with all three exhibiting the same pattern of gene silencing coupled with an enrichment for proteins associated with DNA damage and repair (**Figure 6D**). Collectively, these results show that while both 0N4R and 2N4R isoforms of hTau^P301L^ interact with overlapping proteins, 0N4R and 2N4R isoforms of hTau^A152T^ and hTau^R406W^ interact with distinct target proteins at axonal target sites.

### Phosphorylated hTau is enriched in the somatodendritic compartment

Phosphorylation is critical to regulating hTau’s cellular functions and intracellular localization ^58,22,59^, so we enriched for phosphopeptides and tested whether our hTau isoforms and mutation pairings had different patterns of phosphorylation. We detected several previously identified phosphorylation sites with the majority occurring in the proline-rich region and C-terminus (**Figure S4A**). With a few exceptions, the samples clustered foremost by anatomical region (**Figure S4B**), with several phosphorylated residues reaching significance from PL but none from BLA (**Figure S4C; Table S7**). This is consistent with past work showing phosphorylated tau largely mislocalizes to the somatodendritic compartment^69^ and with the increased AT8 labeling across each mutation in PL neurons (**Figure 1D**). Each of the identified phosphorylated residues has been previously linked to disease pathology^22^. Three phosphorylated residues (pS199, pS202, and pT231) were found at a greater abundance in mutant hTau protein compared to hTau^WT^ and each of them occurred in the 0N4R isoform (**Figure S4D**). pS199 and pS202 were significantly increased in hTau^A152T^, while T231 in hTau^P301L^ was significantly increased compared to hTau^WT^. We then carried out an over-representation enrichment analysis to examine the proteins interacting with these differentially phosphorylated hTau proteins. We restricted the analysis to the three 0N4R hTau proteins (A152T, P301L, and WT) that exhibited significant changes in their phosphorylation pattern. All three interactomes were enriched with synaptic proteins (**Figure S4E**), further connecting the phosphorylated hTau proteins to changes in the dendrites of the cells expressing the proteins.

### Altered mitochondrial function in human iNeurons expressing hTau

Both the 0N4R and 2N4R isoforms of hTau^R406W^ had increased interactions with mitochondrial proteins and GO terms strongly associated with cellular metabolism/aerobic respiration. Although mitochondrial-associated proteins were found to bind each construct of hTau, their enrichment was highly selective for R406W, which argued against a nonspecific interaction during the immunoprecipitation process in which cells are lysed. We sought to functionally validate this interaction in vitro and extend our work to a human cellular context by using human inducible pluripotent stem cells. We engineered the well-characterized AAVS TO-NGN2 KI/WT cell line^70^ (referred to as “control iNeurons”) to express 0N4R hTauWT or 0N4R hTau^R406W^ and used doxycycline-inducible Ngn2 expression to differentiate human iPSCs into glutamatergic neurons (iNeurons) using a previously published protocol^71^ (**Figure 7A**; see Methods). We chose to focus on the 0N4R isoform as it had the strongest mitochondrial signature.

**Figure 7.**
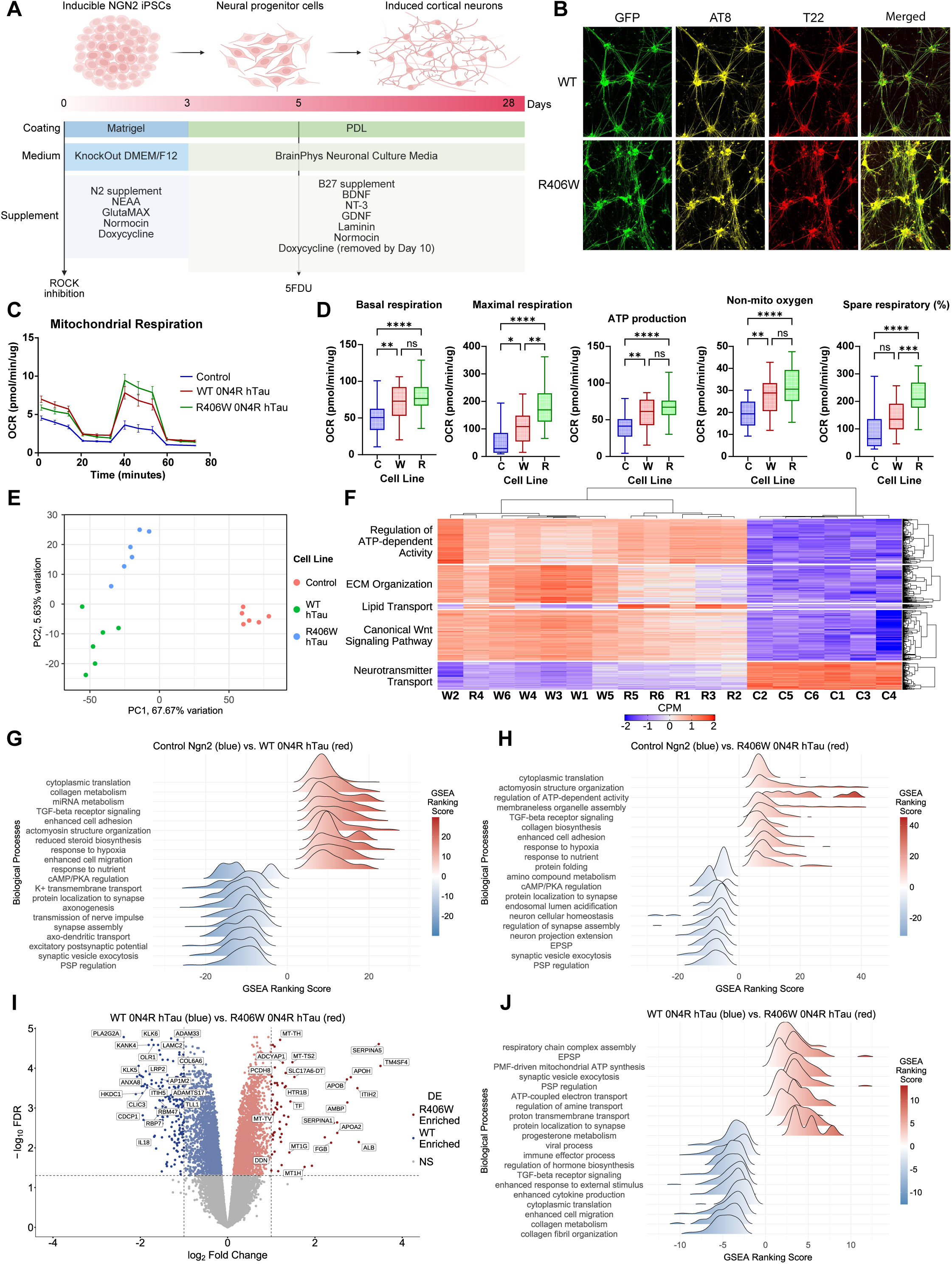
Altered cellular energetics of hTau-expressing iNeurons. **A)** Validated protocol for the differentiation of Ngn2-iPSCs into induced cortical neurons. **B)** D14 WT and R406W iNeurons expressing EGFP (green) were stained with antibodies for AT8 (yellow) or T22 (red). **C)** Oxygen consumption rate (OCR) over time for D28 iNeurons. **D)** Box plots depicting several key metrics (basal respiration, maximal respiration, ATP production, non-mitochondrial oxygen, and spare respiratory % for the control iNeuron (C), WT iNeuron (W), or R406W (R) iNeuron lines. **E**) Principal component analysis (PCA) reveals that the majority (67.67%) of the variance in the transcriptomes of the cell lines is explained by the first principal component followed by only 5.63% for the second principal component. **F)** Hierarchical clustering of the samples based on gene expression. Color scale indicates the level of gene expression from high (red) to low (blue). Genes were split into five major clusters, and an over-representation analysis was run for each cluster resulting in five overarching biological processes. **G-H)** Ridge plot highlights top biological processes enriched for in gene set enrichment analyses (p value > 0.05) carried out between control iNeurons and WT (G) or control and R406W (H). **I**) Volcano plot shows the differentially expressed genes in R406W hTau cells (red) compared to WT hTau cells (blue) (|log2 fold change| > 1 and p-value < 0.05). **J)** Ridge plot highlights top biological processes enriched for in the gene set enrichment analysis (p value > 0.05) carried out between WT and R406W hTau cells.

By 14 days, the iNeurons expressed distinct neuronal markers and exhibit the expected morphological features (**Figure 7B**; **Figure S7A-S7B**). After 28 days, the mitochondrial function of the iNeurons were assayed using a Seahorse XF Analyzer. The oxygen consumption rate (OCR) was remarkably increased in the two hTau cell lines (**Figure 7C**). Both hTau-expressing neuronal lines had elevated levels of basal respiration and ATP production indicating a higher baseline energy demand than the control iNeurons (**Figure 7D**). Interestingly, these cells also exhibited an increase in non-mitochondrial oxygen consumption, a metric that often increases with cellular stress. Relative to control and WT iNeurons, R406W iNeurons demonstrated significant increases in their capacity to respond to the larger energy demand as both the maximum respiration and spare respiratory capacity was significantly elevated compared to both WT and the control iNeurons. WT and R406W iNeurons differed in the extracellular acidification rate (ECAR; **Figure S7C**), where both cell lines were significantly elevated compared to control iNeurons. WT hTau cells overall had significantly higher levels than both R406W hTau and the control iNeurons, which could result from increases in glycolysis or the TCA cycle. These results are reminiscent of those found with the V337M mutation from patient cells^59^, in which the heterozygous and homozygous conditions were each associated with increased basal and maximal rates of respiration. Collectively, these results show that mitochondrial respiration in iNeurons is altered by the expression of hTau.

### hTauR406W alters transcriptional signatures of mitochondrial function in human iNeurons

Given the dramatic shifts in energy production exhibited by the hTau-expressing cells, we were interested in exploring if this corresponded with shifts in the regulation of transcriptional pathways across the three cell lines (**Figure S7D-S7E**). There was distinct separation between cell lines in a PCA analysis with the two hTau cell lines sharing more in common than the control iNeurons (**Figure 7E**), which was also seen with hierarchical clustering of the samples based on gene expression (apart from a single R406W sample (**Figure 7F**). Broadly, the differentially expressed genes (DEGs) largely belonged to five biological processes for the hTau expressing cells: an upregulation of the “regulation of ATP-dependent activity”, “canonical wnt signaling pathways”, “ECM organization,” and “lipid transport”, and downregulation of “neurotransmitter transport” (**Table S8**). Upregulation of ECM organization fits with our proteomic results; although the strongest enrichment was seen with A152T and P301L, there was a significant number of proteins associated with the extracellular matrix in the interactomes of the 0N4R hTau^WT^ and hTau^R406W^ proteins as well.

To further explore how the expression of hTau is affecting the function of these cells, we carried out a gene set enrichment analysis. Terms associated with synaptic function were largely downregulated in both hTau cell lines compared to control iNeurons (**Figure 7G-H**). Several of the significantly enriched processes in the hTau expressing cells, including cytoplasmic translation and cellular response to oxygen and nutrients, further emphasize the metabolic differences from control iNeurons, which was more apparent with R406W than WT. When comparing the R406W iNeurons to the WT iNeurons, we found an upregulation of processes associated with aerobic respiration (**Figure 7I-J**), similar to those found in the binding partners of 0N4R hTau^R406W^ in PL. Interestingly, there are several pathways associated with synaptic firing that are upregulated, suggesting that its synaptic function may fall somewhere in between the control cells and WT. Surprisingly, the pathways that were upregulated in WT and downregulated in R406W also reflect what was seen in the proteomics with 0N4R hTau^WT^, including an increase in terms related to the immune system. Collectively, the transcriptomes of hTau-expressing iNeurons reveal cellular changes parallel to those found from protein-protein interactions, demonstrating that the pathways identified in mouse neurons in vivo underlie distinct mechanisms of specific disease-relevant hTau expression in human neurons.

## Discussion

Here, we leveraged a viral expression strategy to understand how different disease-relevant hTau isoforms and mutations alter synaptic connectivity and hTau functional interactions in excitatory prefrontal cortical neurons. We showed that diverse hTau isoforms and mutations can drive both common and divergent cellular alterations and have provided a resource that can be employed to further explore these changes in a controlled and comprehensive manner. Along with the viral reagents, this resource includes synaptic reconstructions, electrophysiology recordings, multi-site proteomics data, and in vitro functional and transcriptomic changes of human-derived differentiated iNeurons. Expression of each isoform/mutant in mouse PL-to-BLA excitatory pyramidal neurons produced remarkably similar patterns of synaptic loss and spine neck remodeling, suggesting a highly conserved response to disease-relevant mutant hTau. Despite this consistency, analyses of the intracellular protein-protein interactions revealed stringent specificity for each isoform, mutation, and subcellular location. Our results highlight a prominent role for hTau^R406W^ in mitochondrial and energy-related processes, and we confirmed this functionality in mitochondrial assays from hTau^WT^- and hTau^R406W^-expressing human Ngn2-induced neurons *in vitro*. Collectively, these results show a common implementation of synaptic disconnection across hTau mutants masks significant subcellular molecular heterogeneity of protein-protein interactions.

Compared to transgenic mouse lines, the viral approach used here has several advantages. Targeted brain injection strategies allow direct control over the spatial and temporal expression patterns, and our Cre-dependent construct design allows cell-type specificity when combined with either Cre-expressing viruses or Cre knockin driver lines. The viral strategy also allows direct comparisons across different isoforms and mutations all on a common genetic background with rigorous comparison to matched hTau^WT^-expressing controls; such an approach is not possible with the random insertion of hTau constructs and not yet feasible with humanized knockin mice (which are not restricted to specific isoforms; see^72-74^). Although these strategies increase construct and face validity, they preclude interrogation of the effect of expression on specific cell-types or at subcellular locations. On the other hand, viral overexpression of hTau constructs does not recapitulate outright tau tangle neuropathology^21^, limiting the relevance of our results to relatively early disease phases. Additionally, viral strategies produce payload overexpression, and viral load on its own might drive artificial activation of the immune system that likely influences the underlying biology^75^. Carrying out the experiments in a standard mouse strain similarly allows us to better compare our results to popular tauopathy models (AAVs, transgenic mice) where endogenous tau is present, but it also excludes us from attributing all of the observed changes solely to the presence of the humanized tau protein.

Here, we expressed these constructs in mouse PL cortex, which is structurally and functionally organized similarly to the human regions sensitive to many primary tauopathies^76^. The observed synapse loss on PL-to-BLA pyramidal neurons is consistent with prior work suggesting large-scale synapses disconnection within associative circuits prior to observable tangle pathology^32,33^. Synapse loss seen on the proximal and main apical branches was largely absent from the distal tuft dendrites, which may reflect the distinctly different patterns of afferent excitatory and inhibitory synaptic connectivity on these branches^77-80^. Independent of whether the maintenance of excitatory connections on distal dendrites results from a distance-dependent intracellular gradient of hTau molecular function or from circuit-specific patterns of local activity generated *in vivo*, the selective preservation of some inputs at the expense of others will ultimately distort multi-compartment forms of synaptic integration that are typically associated with pyramidal cell computations^81^. At a finer scale, the increased distances between synapses and the reduction of spine neck lengths should synergize to decrease the engagement of nonlinear dendritic mechanisms in response to synaptic input. These changes, coupled with the observed uniform hTau-mediated reduction of spike output form somatic current injections, strongly argue that different disease-related hTau mutants alter the input-output relationship in prefrontal cortical neurons in a similar way.

Surprisingly, the similarity of these changes across the hTau isoforms and mutations contrasted with significant specificity within their protein-protein interactions. The combination of cell-type specific expression, multisite microdissections, and high-sensitivity mass spectrometry allowed intracellular hTau protein-protein interactions to be analyzed separately for somatodendritic and distal axonal sites of PL excitatory neurons. We found three specific signatures: one in which somatodendritic 0N4R hTau^A152T^ and 0N4R hTau^P301L^ interacted with a large number of proteins related to the extracellular matrix; a second in which somatodendritic 0N4R and axonal 2N4R hTau^R406W^ interacted with mitochondrial proteins associated with cellular respiration and energy metabolism; and a third in which a number of axonal mutant hTau proteins, including both isoforms of hTau^P301L^, engaged with proteins associated with chromatin accessibility, nuclear gene repression, and DNA repair. Although hTau has known interactions with ECM proteins^82,83^, mitochondrial proteins^59^ and nucleoproteins^57,84^, the isoform-, mutation-, and subcellular specificity underlying these interactions has remained unidentified. Presumably, the interaction with nuclear proteins isolated from PL axons in the BLA site reflects transcellular transport of the hTau protein. Although we could not visualize FLAG^+^ proteins in the nucleus of BLA neurons, transcellular propagation of hTau protein has been shown in hippocampal and cortical circuits with very similar viral constructs (though see^21,68^). The notion that these results extend beyond the mouse PL cortex and to human cellular contexts is strengthened by the observation that hTau^R406W^ expression in human cells produced changes in mitochondrial function and dysregulation of transcripts related to cellular metabolism.

Our functional experiments show that expression of both hTau^WT^ and hTau^R406W^ dramatically altered mitochondrial function in human differentiated neurons in vitro. These results are consistent with those obtained from the V337M mutation in differentiated iPSCs^59^, and provide strong evidence that hTau imparts strong control over cellular respiration and mitochondrial function. Our proteomics and transcriptomics results suggest this interaction is implemented by both direct binding between hTau and mitochondrial-associated proteins, and by transcriptional modulation of genes associated with cellular respiration. Using MitoCharta 3.0, we detected 123 mitochondrial proteins in the combined PL and BLA dataset (**Figure S8A**). Unsurprisingly, the majority were bound by hTau^R406W^ (88 of 123) and hTau^WT^ (77 of 123), substantially more than hTau^A152T^ (42 of 123) and hTau^P301L^ proteins (48 of 123). The proteins largely localize to the mitochondrial inner membrane, the location of the major electron transport complexes (ETC) complexes and the translocases Timm9 and Timm10, and to the mitochondrial matrix (**Figure S8B**). Transcriptionally, the genes upregulated in the R406W iNeurons compared to WT iNeurons suggest enrichment in pathways involving the assembly of the ECM complexes, its subunits, and mitochondrial translation (**Figure S8C).** This connects nicely with what was observed in the proteomics, as well as suggests potential mechanisms, including rearrangement of ETC complex composition and altered protein synthesis, by which these cells could increase their metabolic capabilities. Future work should define exactly how these mechanisms interact to shape mitochondrial function in vivo over time and in different cell types.

Previous work has demonstrated different tau mutations engage distinct mechanisms promoting circuit and cellular dysfunction, but the comparisons are necessarily made across studies with different endpoints (e.g., aggregation, LTP protocols, or mouse behaviors), often from mice created with different transgenic insertion strategies expressing mutations on different hTau isoforms. By contrast, our tools can be delivered to specific cell types and with rigorous hTauWT-expressing controls. As a resource, our datasets can be leveraged to generate novel hypotheses about cellular pathways engaged by disease-relevant mutant hTau. The diversity of our results, even when restricted to PL cortex, suggest there will be circuit- and cell-type– specific rules by which hTau proteins interact with their cellular targets. Moreover, the reagents made here can be leveraged in cellular systems and delivered to mouse or rat Cre driver lines to test the mechanisms of selective cellular vulnerability in different circuits and cell types, or to drive mutant hTau in mice with mixed pathologies related to FTD or AD.

Our results have potential implications for strategies aimed at precision therapeutic approaches for tauopathies. Recent strategies focused on increasing the resilience of synaptic connections by manipulating neurotrophin signaling^85,86^ may limit synapse loss, normalize neural computations, and ultimately restore information processing in cortical circuits. Optimistically, the uniform synaptic changes seen here across the hTau isoform/mutation combinations suggest these efforts might have utility for individuals harboring a broad swath of tau mutations, including those not examined here. At the same time, our results also suggest the success of such an approach might be ultimately limited by the specific alterations in tau interactions inherent to each isoform and mutation. Future work should leverage our approach to reverse engineer the molecular mechanisms of selective circuit vulnerability to mutant hTau. This should include driving hTau expression in nonneuronal cell types (e.g., astrocytes and oligodendrocytes) using viral tools now available to gain selective genetic access to these cell types^87,88^.

## Lead Contact

Further information and requests for resources and reagents should be directed to Erik Bloss.

## Materials Availability

All plasmids for the viral constructs generated here will be deposited at Addgene.

## Data Code and Availability

All datasets and analysis code will be deposited at GitHub and at a publicly available repository upon publication.

## Acknowledgements

We thank all members of the Bloss laboratory for helpful discussion and comments; Drs. P. West, G. Howell, K.M.S. O’Connell, and R. Burgess for experimental and editorial suggestions; P. Heinrich, P. Finger, and the JAX Microscopy and Electron Microscopy Cores for assisting with experiments; and the Animal Care team at JAX. This work was supported by funding from The Jackson Laboratory’ Director’s Innovation Fund (to E.B.B.), by T32AG062409 (to E.N), and by F31AG094270 (to E.N.). E.B.B. is supported by the National Institutes of Health (R01AG079877).

## Author Contributions

Conceptualization, E.N. and E.B.; methodology, E.N. J.P-T., S.P., C.R., T.Z., T.S., B.H., R.M., and E.B.B.; formal analysis, E.N., J.P-T., T.Z., T.S., B.H., R.M., and E.B.B.; investigation, E.N., J.P-T., T.Z., T.S., B.H., R.M., and E.B.B.; writing – original draft, E.N. and E.B.B.; writing – reviewing & editing, E.N. and E.B.B.; visualization, E.N. and E.B.B.; supervision, E.B.B.; funding acquisition, E.N. and E.B.B.

## Materials and Methods

### 1. Experimental model and subject details

#### Mice

All C57BL/6J mice (JAX stock #000664) were bred and group housed at the Jackson Laboratory with a light/dark cycle of 12 hr. and fed a standard extruded chow diet (LabDiet #5K52). Mice of both sexes between the ages of 4 to 6 months old were used in the experiments. Littermates were randomly assigned to experimental groups.

#### Cell Culture

The KOLF2.1J AAVS1-TO-NGN2 KI/WT iPSC line (JAX #JIPSC002070), a doxycycline-inducible cell line that utilizes NGN2 expression to rapidly differentiate cells into cortical excitatory neurons, was used as the parent line to create the two hTau cell lines. PiggyBac plasmids encoding wildtype or R406W versions of EGFP-T2A-TAU, along with a plasmid encoding PiggyBac transposase, were introduced to the KOLF2.1J cells using nucleofection (Lonza 4D nucleofector) according to kit manufacturer’s instructions (Mirus Bio). KOLF2.1J cells were maintained on recombinant vitronectin coated plates in StemFlex media (both from Life Technologies). After 24 hours, puromycin (Invivogen) was added to 1 ug/ml final concentration. Cells were maintained for several passages in puromycin to enable stable cell selection. Selected cells were maintained as a pool prior to downstream experimental procedures.

Cells were maintained in a cell culture incubator at 5% CO_2_ at 37°C. The three iPSC lines were grown in StemFlex media (Gibco, cat #A3349401) in six-well plates coated with Synthemax (0.025 mg/mL, Corning, cat #3535) or Vitronectin (0.01 mg/mL, StemCell Technologies, cat #07180). ReLeSR (StemCell Technologies, cat# 100-0483) was used to dissociate the cells during passaging. To generate neural progenitor cells (NPCs), iPSCs were dissociated using Accutase (StemCell Technologies, cat # 07922) and plate onto six-well plates coated previously with Matrigel (1 mL/well, Corning, cat #354277) at a density of 1 – 2 x 10^6^ cells/wells in induction media, consisting of KnockOut DMEM/F-12 media (Gibco, cat #12660012) supplemented with N2 supplement (100X, Gibco, cat #17502048), non-essential amino acids (100X, Gibco, cat #11140050), Gluta-MAX (100X, Gibco, cat #35050061), Normocin (50X, Invivogen, cat#NC2209896), and doxycycline (2 mg/mL, Sigma, cat #D9891). Rock inhibitor (100X, Selleck Chem, cat #S1049) was only included for the first day. Media was replaced daily and after three days the cells were dissociated with Accutase and plated onto plates coated with PDL (0.1mg/mL, Sigma, cat# P7886M) or PLO (0.1mg/mL, Sigma, cat #P3655). Cells were maintained in BrainPhys neuronal media (Stem Cell, cat #05790) containing B27 supplement (50X, Gibco, cat #A35828), BDNF (10 ng/mL, Peprotech, cat #PHC7074), NT-3 (10 ng/mL, Peprotech, cat #450-03-10UG), GDNF (10 ng/mL, Peprotech, cat #450-10-10UG), laminin (1 ug/mL, R&D system, cat #3446-005-01), Normocin (50X), and doxycycline (2 mg/mL). Following this, half of the media was replaced three times a week. 5FDU was added between day 5 and 6 and doxycycline was removed no later than day 10.

### 2. Method Details AAV Production

To facilitate creation of AAV.hSYN.FLEX plasmids encoding EGFP-2A-TAU, synthetic DNA fragments encoding EGFP-T2A-TAU (2N4R and 0N4R variants) were synthesized by Twist Bioscience (South San Francisco, CA). The open reading frame of iGluSnFR3.v857.SGZ.codonopt was removed from plasmid pAAV.hSyn.FLEX.iGluSnFR3.v857.SGZ.codonopt (Addgene #175182), and the synthetic EGFP-T2A-TAU fragments were inserted in its place. Additional fragments encoding P301L, R406W, and A152T variants were also synthesized at Twist Bio and were used to replace the wildtype sequences by traditional restriction digestion-ligation cloning. All plasmid production was done in NEB Stable *E. coli* (New England Biolabs, Beverly, MA) at 30°C.

To create ‘floxed’ versions of hSYN.EGFP-T2A-TAU plasmids, *in vitro* CRE-recombinase reactions (New England Biolabs, Beverly, MA) were performed using the hSYN.FLEX.EGFP-T2A-EGFP plasmids with manufacturer’s recommended protocols. Resulting plasmids were transformed into NEB Stable *E. coli* and resulting colonies were screened by PCR for correct orientation and further characterized by sequencing to confirm correct LoxP/Lox2272 recombination.

AAV2/1 particles were produced by triple-transfection in AAV-293 HEK cells (Agilent) using JetOptimus (Polyplus, France). pAAV2/1 (Addgene plasmid #112862) and pAdDeltaF6 (Addgene plasmid #112867) plasmids were obtained from UPenn Vector Core (Philadelphia, PA). In some instances, helper plasmid pALD-HELP (Aldevron) was used instead of pAdDeltaF6 while keeping molar ratios during transfection constant. Following transfection, media was changed to low-serum containing media and was collected at 48 hrs. Media was replaced and cells and media were collected together at 72 hrs. Cells were pelleted and subjected to multiple freeze-thaw cycles in ‘Gradient buffer’ (10 mM Tris-HCl pH7.6, 150 mM NaCl, 10 mM MgCl2). Following treatment with benzonase (E1014, Millipore Sigma) for 30 min at 37°C, AAV particles in clarified supernatant were precipitated by addition of PEG-8000/NaCl to a final concentration of 8% PEG-8000/0.5M NaCl, incubated on ice for 1 hr., and pelleted at 3500xg. Precipitated AAVs were resuspended in gradient buffer and combined with cellular lysate. After clarification, AAV solutions were loaded onto iodixanol gradients and centrifuged in a Thermo F50L rotor @ 48,000 RPM for 2 hrs at 18C. AAVs were recovered from the 40% iodixanol gradient. Buffer exchange (to 1X Hanks buffered saline/0.001% Pluronic F-68) and concentration was performed using Amicon 100 kD cutoff centrifugal filters with three buffer exchanges. Concentrated viruses were aliquoted to low-binding tubes and stored at -80C. Titration was performed by dPCR (Qiagen) using primer/probe against ITRs.

#### Intracranial Viral Injections

Viral constructs were stereotactically injected into the prelimbic cortex (+1.95 mm, 0.80 mm, and 1.75 mm with the mouse head tilted at a 15-degree angle) or into the basolateral amygdala (-1.6 mm, 3.3 mm, and 4.2 mm, with all coordinates reported as mm anterior/posterior from Bregma, lateral to the midline, and ventral from the surface of the brain). All surgeries were performed in 4 to 6-month-old C57BL/6J mice under isoflurane anesthesia (1-3%) using a Kopf stereotactic apparatus (#1902) equipped with a Zeiss microscope (V.20). At each site, 100 nL of viral suspension was injected over a period of three minutes using glass pipettes (∼30 µm diameter tips) with Drummond Nanoinjectors driven by a micromanipulator (Sutter MP-285). Injection pipettes were left in place for 4 to 5 minutes and then slowly retracted. To express hTau in PL-to-BLA neurons, mice received injections of the viral hTau constructs in PL and a retro-Cre construct (Addgene #55636) in BLA. The dendritic spine analysis leveraged a within-animal comparison in which Cre-dependent EGFP was injected into left PL, Cre-dependent hTau injected into right PL, and a retro-Cre construct bilaterally into BLA. For the mass spectrometry experiments, a 1:1 viral mixture of the hTau expressing viral construct and CaMKII-Cre (Addgene #105558) was injected into the prelimbic cortex. Mice were given carprofen and Ethiqa to aid in recovery and monitored for 5 consecutive days to ensure proper wound healing.

#### Immunofluorescent Labeling and Confocal Microscopy

Mice were anesthetized with Tribromoethanol (800 mg/kg, i.p.) and transcardially perfused with ice-cold 4% depolymerized paraformaldehyde (EMS) in 0.1 M PB. For immunofluorescent protein labeling, coronal slices were cut on a Vibratome (50 µm-thick) and were incubated in blocking buffer containing 2% BSA and 0.3% Triton X-100 followed by incubation in primary antibodies: FLAG (1:10,000, rabbit, Sigma, cat# F7425), AT8 (1:1000, mouse, ThermoFisher, cat #MN1020), IBA-1 (1:1000, rabbit, Fujifilm, cat#019-19741), CLEC7A/mDectin-1 (1:100, Invivogen, rat, cat#NC2684392), and GFAP (1:1000, chicken, Origene, cat# TA309150). For immunofluorescent imaging of spines, slices were incubated in blocking buffer containing 2% BSA, 1% fish gelatin, 5% goat serum, and 0.3% Triton X-100 followed by incubation in anti-FLAG primary antibody (1:10,000, mouse, cat# F1804). EGFP signals were not enhanced with antibodies. Slices were washed in buffer containing 0.3% Triton X-100 and followed by incubation of the fluorescent-conjugated secondary antibodies diluted in the same blocking buffer. Slices were mounted onto microscope slides and coverslipped with Vectashield.

For postembed immunogold labeling (Figure 1), mice were perfused with 4% paraformaldehyde + 0.125% glutaraldehyde and tissue punches (3 mm diameter) were rapidly vitrified with a Leica ICE high-pressure freezer. Tissue blocks were freeze-substituted, embedded into HM20 resin, polymerized by UV light, cut (70 nm-thick) on a ultramicrotome, and collected onto carbon-coated Formvar slot Grids (EMS). Postembed immunolabeling was performed as previously described^89^. Grids were imaged with a JEOL 1230 transmission electron microscope and images were collected with a CMOS camera (Hamamatsu).

Confocal microscopy for dendritic and axonal analysis was performed using a Leica Confocal Microscope (SP8) equipped with multiple objectives, laser lines, and PMT/HyD photodetectors. For AT8/EGFP and IBA1/CLEC7A analyses, tiled z-stacks were collected with a 40x 1.3 NA objective, and fluorescent intensity of individual cellular regions of interest were measured for all of the channels. Dendritic branches and spines were imaged with a 63x 1.4 NA objective, at 0.05 x 0.05 x 0.1 µm voxel sizes, and processed with Lightning Deconvolution. Dendritic spines and axons were analyzed in NeuronStudio^90^, which allows a semiautomated analysis of branch characteristics and individual spine head volumes and neck lengths.

#### Electrophysiology

Mice expressing EGFP (control) or the 0N4R form of hTau^WT^, hTau^A152T^, hTau^P301L^, or hTau^R406W^ were anesthetized with isoflurane before being sacrificed for electrophysiology recordings. Brains were rapidly removed and placed in ice-cold cutting artificial CSF (aCSF) containing (in mM): 93 NMDG, 2.5 KCl, 1.4 NaH_2_PO_4_, 30 NaHCO_3_, 20 HEPES, 25 D-Glucose, 3 Myoinositol, 5 Sodium L-Ascorbate, 2 Thiourea, 6.1 NAC, 3 Sodium Pyruvate, 0.01 Taurine, 10 MgSO_4_.7H2O, 0.5 CaCl_2_.2H_2_O, pH adjust to 7.4 and osmolarity to 300 mOsm. Acute transverse slices (250 µm) were cut with a vibratome (Leica, VT1000S) in ice cold cutting aCSF. Slices were incubated at 34℃ in carbogen bubbled cutting aCSF for 17 minutes. Slices were held at room temperature for 1-4 hours before recording in holding aCSF containing (in mM): 94 NaCl, 2.5 KCl, 1.4 NaH_2_PO4, 25 NaHCO_3_, 20 HEPES, 25 D-Glucose, 3 Myoinositol, 2 Thiourea, 5 Sodium L-Ascorbate, 6.1 NAC, 3 Sodium Pyruvate, 0.01 Taurine, 2 MgSO_4_.7H_2_O, 2 CaCl_2_.2H_2_O, pH adjust to 7.4 and osmolarity to 300mOsm. Whole-cell current clamp recordings were made from EGFP-expressing PL neurons at 34℃ under visual guidance of a video microscope (Olympus, Moment CMOS camera) using patch pipettes with resistance of 3-5 MΩ filled with potassium gluconate based internal solution (in mM): 126 K-Gluconate, 4 KCl, 10 HEPES, 4 MgATP, and 0.3 NaGTP, 10 Na-phosphocreatine, 0.2% biocytin, pH adjust to 7.4 and osmolarity to 290 mOsm. Recordings were performed in an aCSF solution containing (in mM) at 34℃: 119 NaCl, 2.5 KCl, 2.5 CaCl.2H_2_O, 1 MgSO_4_, 1.25 NaH_2_PO_4_, 23 NaHCO_3_, 10 D-Glucose. Recordings were acquired at a 10 kHz sampling frequency and digitized at 2 kHz in current clamp mode. Series resistance and capacitance were monitored and compensated throughout recordings. Neurons with >40 MΩ series resistance were excluded. Changes in input resistance and spike output were measured by injecting 5s square step current into the soma ranging from 0 pA to +150 pA in 50 pA intervals. Rheobase was measured by injecting 5s-long depolarizing ramp stimulus (0-300 pA) current step into the soma; input resistance, inter-spike-interval, and rheobase were calculated in Easy Electrophysiology.

#### Mass Spectrometry Data Acquisition

6-month-old C57BL/6J mice received bilateral injections of one of eight different hTau constructs (0N4R hTau^WT^, 2N4R hTau^WT^, 0N4R hTau^P301L^, 2N4R hTau^P301L^, 0N4R hTau^A152T^, 2N4R hTau^A152T^, 0N4R hTau^R406W^, or 2N4R hTau^R406W^) in PL. After three months of expression, mice were anesthetized with Tribromoethanol (800 mg/kg, i.p.) and transcardially perfused with ice-cold 1X PBS. PL and BLA were microdissected from the tissue, snap-frozen in liquid nitrogen, and stored at -80°C.

Each tissue sample was transferred to a thick-wall 2 mL microcentrifuge tube and had 250 μL of lysis buffer added (20 mM HEPES, pH 7.4; 10 mM KCl, 0.05% w/v NP-40, 1X Roche protease/phosphatase inhibitors). A 5 mM steel bead (Qiagen) was then added to each sample tube, and samples were lysed in the Tissue Lyser II (Qiagen) for 2 minutes at 30 oscillations per second. All samples were then waterbath sonicated at 37Hz for 5 minutes (30 seconds on/30 seconds off). Following the lysis steps, samples were centrifuged at 2000 x g for 15 minutes at 4°C to clarify the lysate by removing heavy debris. Lysate supernatant was transferred to a new 1.5 mL microfuge tube, and the protein concentration was determined using a MicroBCA quantification assay according to manufacturer protocol (Pierce).

Immunoprecipitation was performed using the Monoclonal anti-Flag M2 Antibody produced in mouse (F1804-200UG; Sigma) bound to M-280 Sheep Anti-Mouse IgG Dynabeads™ (11-202-D; Fisher Scientific) according to manufacturer protocols (5 μL of antibody per 5e10^7^ beads for each sample). The antibodies were bound to the beads rotating end-over-end at 4°C for 16 hours and washed 3 times in PBS containing 5 mg/mL bovine serum albumin on a magnet; following the last wash beads were brought up in 100 μL of Binding/Wash Buffer A (20 mM HEPES, pH 7.4; 10 mM KCl; 50 mM NaCl, 1 mM MgCl_2_; 0.05% w/v NP-40) for each sample. The bound beads were added to 250 μg of lysate diluted to 100 μL in Binding/Wash Buffer A (200 μL total). Additionally, both IgG control samples (beads with no anti-Flag antibody) and Mock control samples (beads with anti-Flag antibody but incubated with non-Flag lysates) were added to the study. The sample and antibody beads were then incubated at 37°C for 30 minutes in a Thermomixer with agitation (700 rpm). Bound beads were then washed four times on a magnetic rack with 1 mL of Wash Buffer A, then two times with Wash Buffer B (20 mM HEPES, pH 7.4; 10 mM KCl; 0.05% w/v NP-40). At each wash tubes were vortexed at max speed for 15 seconds. Following the last wash, the final wash supernatant was removed from the tubes in the magnetic rack and 50 μL of 50 mM HEPES, pH 8 was added to each sample. Protein bound bead samples were then reduced with 10 mM dithiothreitol for 30 minutes at 42°C, alkylated with 15 mM iodoacetamide for 20 minutes at 25°C in the dark, and digested with 500 ng of sequence grade trypsin (Promega) at 37°C for 20 hours. All on-bead digest steps were performed in a Thermomixer with agitation (700 rpm) to keep the beads suspended. Supernatant (peptide containing) was removed from each digest on a magnetic rack and transferred to a new tube containing 1 μL of 10% TFA to quench the samples (checked pH to ensure it was pH 3-4 or adjusted accordingly). Peptide samples were then purified using Millipore C18 zip-tips (Millipore, ZTC18S096) according to manufacturer protocol and dried using a vacuum centrifuge.

A separate subset of peptides from the samples, following immunoprecipitation also underwent a phospho-enrichment. Dried peptide samples from C18 zip-tip purification were resuspended in 50 μL of reconstitution solution (5% ACN, 50 mM ammonium bicarbonate) by pipetting a minimum of ten times, followed by vortexing. The samples then had 5 µL of 5% TFA added to mimic the steps as if a secondary digest was performed. From there the phospho-enrichment was completed per the PTMScan® IMAC Fe-NTA Magnetic Beads manufacturer protocol with volume adjustments to fit the starting sample volume while keeping ratios. The PTMScan® IMAC Fe-NTA Magnetic beads were resuspended by rotating end-over-end on a rotator 10 minutes at room temperature; 20 µL of 25% bead slurry (5 µL packed beads) were used per sample and transfer to a 2.0 mL microcentrifuge tube. The IMAC beads were washed three times using a magnetic cassette with 1 mL Wash Buffer (0.1% TFA, 80% acetonitrile). After the last bead wash, the reconstituted samples were transferred to the tube with the IMAC resin and mixed by pipetting, then incubated on a Thermomixer for 30 minutes at room temperature with agitation (700 rpm) to keep the beads suspended. After the incubation, the supernatant (phospho-depleted solution) was separated from the beads (bound phospho-peptides), and the beads were washed three times with 1 mL of Wash Buffer. Phospho-peptides were then eluted from the beads by adding 50 µL of Elution Buffer (50% acetonitrile, 2.5% ammonia) and incubated on the Thermomixer for 5 minutes at room temperature with agitation (700 rpm). The beads were allowed to settle on a magnetic rack and the supernatant containing the eluted phospho-peptides was transferred to a clean 1.5 mL microcentrifuge tube. The elution process was repeated one more time, the supernatants were combined, and 40 µL of 20% TFA was added to acidify the eluate. The eluted phospho-peptides were dried in a vacuum centrifuge and reconstituted in 40 μL of 0.1% TFA and zip-tipped according to manufacturer protocol (slight modification - use a 40% acetonitrile with 0.1% TFA elution). The samples were then dried in a vacuum centrifuge, reconstituted in 20 μL of 0.1% TFA, and transferred to a mass spectrometry vial for analysis.

Dried peptide samples were reconstituted in 20 µL of 98% H2O/2% ACN with 0.1% formic acid by pipetting and vortexing. Samples were then analyzed using data independent acquisition mass spectrometry analysis (DIA-MS) on a Thermo Orbitrap Astral mass spectrometer coupled to the Vanquish Neo liquid chromatography system in the Mass Spectrometry and Protein Chemistry Service at The Jackson Laboratory as described previously^91^. An EASY-Spray column (PepMap Neo C18, 75 µm x 150 mm, #ES75150PN) was utilized for chromatography separation of the peptides. Buffer A (100% H2O with 0.1% formic acid) and Buffer B (100% acetonitrile with 0.1% formic acid) were used for a 25-minute gradient ramping from 98%A/2%B to 10%A/90%B with a flow rate of 600 nL/min as described previously^91^. Detailed mass spectrometry instrument parameters can be found as described previously^91^.

#### Immunocytochemistry

100,000 NPCs were seeded per well onto 8-well chambered coverslips (Ibidi, cat#80826) coated with heavy-weight PDL (0.1 mg/mL, Millipore Sigma, cat #P7405), fibronectin (1:500, Millipore Sigma, cat #FC010) and laminin (1:1000). The following steps took place at room temperature, and each wash step was repeated three times, unless otherwise noted. Cells were washed once with PBS, incubated with 4% paraformaldehyde for 20 minutes, and then washed with PBS. Cells were incubated in blocking buffer for 30 minutes, washed with PBS, and incubated for either 3 hours in primary antibody at room temperature or overnight at 4°C. The primary antibodies included: GFP (1:5000, Abcam, cat # ab13970), AT8 (1:500, mouse, ThermoFisher, cat #MN1020), T22 (1:500, rabbit, Sigma-Aldrich, cat # ABN454), NeuN (1:500, rabbit, Abcam, cat #ab104225), and Phospho-T231 (1:500, Abcam, cat # ab151559). Cells were washed in PBS, incubated in fluorescent-conjugated secondary antibodies for 1 hour, and then washed again in PBS. Each well was filled with PBS, protected from the light, and stored at 4°C. Confocal microscopy was performed using either the microscope detailed above or a Leica Stellaris 8. Tiled z-stacks were collected with a 40X 1.3 NA objective.

#### Seahorse Assay

100,000 NPCs were seeded per well onto a PLO-coated XFe96 Cell Culture Microplate (Agilent, cat #103794-100) and received regular ½ media changes as described above. At 28 days post induction, mitochondrial function was assessed using the Seahorse XFe 96 Flux Analyzer according to the instruction provided in the Seahorse XF Mito Stress Test kit (Agilent, cat #103015-100). The Seahorse Media (Agilent, cat #103335-100) was supplemented with 1 mM pyruvate (Agilent, cat #103578-100), 2 mM glutamine (Agilent, cat #103579-100), and 10 mM glucose (Agilent, cat #103577-100). Following the addition of the seahorse media, bright field images were taken of each well using the Cytation instrument. Empty and sparsely populated wells were excluded from the downstream analysis. The plate remained in the instrument until the required hour of outgassing time had elapsed. At this point, it was transferred to the Seahorse XFe 96 Flux Analyzer, and the assay was run using the following drug concentrations: oligomycin (2.5 µM), FCCP (2 µM), and Rotenone/Antimycin (0.5 µM). Once completed, the plate was briefly centrifuged and washed once with PBS. 50 μL of cell lysis buffer, consisting of 1 protease inhibitor complex tablet (Thermo Scientific, cat #A32961) dissolved in 10 mL of NP-40 lysis buffer (Thermo Scientific, cat #J60766.AK), was added to each well and the plate was incubated on ice for 30 minutes with gentle agitation every 5 minutes. Supernatant was transferred to a new 96-well plate and centrifuged at 14,000 G for 10 minutes. Each sample was diluted 1:10 with 1X PBS and then the protein concentration was assayed using the Micro BCA™ Protein Assay Kit (Thermo Scientific, cat # 23235). Absorbance for each well was recorded at 562 nm using Spectramax i3 and concentrations for each sample were calculated using the standard curve generated from the standards. Total protein was then used for well-well normalization with the seahorse assay using the Wave software.

#### RNA sequencing

200,000 to 300,000 NPCs were seeded per well onto PDL-coated 24-well plates and received regular 1/2 media changes as described above. At 28 days post induction, cells were washed with 1X PBS and lysed in RNA shield. Samples were briefly spun and stored temporarily in 4°C until shipped to Plasmidsaurus for 3’ mRNA-seq. RNA was isolated with a bead-based extraction method. mRNA was then converted into double-stranded cDNA, tagmented and amplified. The library was sequenced on the Illumina NovaSeq X Plus with asymmetric paired-end sequencing.

### 3. Quantification and statistical analysis

#### Immunohistochemistry analysis and dendritic spine quantification

Relationships between EGFP intensity and AT8 intensity were analyzed using Spearman’s correlation (**Table S9**). Dendritic spine densities were calculated as spines/µm, averaged over 5 branches for each mouse, and analyzed as paired two-tailed t-tests between the EGFP and hTau hemispheres. Densities across hTau constructs were analyzed with one-way ANOVAs followed by Tukey/Bonferonni posthoc tests with corrections for multiple comparisons. For the analysis of nearest neighbor (NN) distances, each NN was calculated and their distributions were analyzed with Kologorov-Smirnov tests. The distribution of spine neck lengths and spine head volumes were made on a per-branch basis, averaged across mice, and tested for significance with two-way ANOVAs with bin and construct as main factors; Sidak-Holmes posthoc tests were used with corrections for multiple comparisons.

#### Analysis of the LC-MS/MS data

All Orbitrap Astral RAW data files underwent automatic UniProtKB Mus musculus (sp_tr_incl_isoforms TaxID=10090; v2024-05-01) protein database searching using CHIMERYS with the Inferys 3.0 fragmentation prediction model through the Thermo Ardia Server and connected Proteome Discoverer workstation (Thermo). All unique sequences for the models in the study were also added to the protein databases to ensure all custom sequences were present. All other search parameters were set as described previously^91^ with the addition of a variable phosphorylation modification (+80-Da on Ser/Thr/Tyr amino acids). Sample data was normalized to total ion signal in the samples and protein abundance-based ratio calculations were used for summed abundances, followed by a background-based t-test and Benjamini-Hochberg multiple corrections testing. Protein group master proteins determined by strict parsimony in Proteome Discoverer were utilized. The data was also run with and without imputation by low abundance resampling for comparison. During the comparison, both the IgG and Mock controls were used to perform background subtractions for non-specific binding. Peptides were filtered for using the following criteria: false discovery rate <= 0.05, coefficient of variation % <= 35%, and peptide spectral match >= 10. Keratins, known contaminants in mass spectrometry experiments, were excluded from the downstream analysis.

Analyses were carried out in R (v4.5.0) and RStudio (v2024.12.1) unless otherwise stated. For all analyses, mutant to WT protein comparisons were always carried out within isoform (ex. 0N4R hTau^A152T^ will only be compared to 0N4R hTau^WT^). To be classified as a binding partner of a hTau protein, the log2 abundance ratio had to be significantly higher than that of the combined mock and IgG controls (p < 0.05). To be considered differentially bound by the mutant hTau protein compared to hTau^WT^, it was either 1) enriched in the mutant hTau interactome compared to control (p < 0.05) and enriched for compared to hTau^WT^ (log_2_ abundance ratio >= 1, p < 0.05) or 2) enriched for in the hTau^WT^ protein interactome (p < 0.05) but depleted in the mutant interactome compared to hTau^WT^ (log2 abundance ratio <= -1, p < 0.05). Identified hTau binding partners were compared to a benchmark dataset, which consisted of 14 previously published mass spectrometry datasets that identified the binding partners of tau using a variety of model organisms and methods. Proteins were classified as novel, known, or as a close homology to a known interactor (ex. different subunit). A network map illustrating the known protein-protein interactions between hTau binding partners was generated using the STRING database (v12.0) and Cytoscape (v3.10.4). The proteins largely belonged to 11 different biological categories, and a subset of known and novel binding partners of each category were displayed. Grouped abundances were log2 transformed before being used in the principal component analysis with PCAtools (v2.20.0) and ggplot2. Volcano plots were created using ggplot2 (v4.0.3). P value was scaled using -log10 and proteins were characterized as differentially bound if they passed the significance requirements (vs control: log2(abundance ratio) >= 1, p <0.05; vs WT: log2(abundance ratio) >= 1, p < 0.05 or log2(abundance ratio) <= - 1, p < 0.05). The upset plot was created with the ComplexUpset package (v1.3.3). To create the circular packing graphs, the UnitprotR (v.2.5.1) was used to correlate gene ontology terms with the Uniprot accession ID for each of hTau’s identified binding partners. The specific biological processes and cellular components were grouped into broader categories using MGI GO Slim and the circle packing graphs were created using pack circles (v0.3.7) and ggplot2. The heatmaps were generated from proteins associated with the GO terms indicated using the normalized log2 abundance ratios and ComplexHeatmap (v2.26.1). For all over-representation analyses (ORA), the differentially bound proteins were separated into two groups: up (enriched for in the interactome) or down (depleted for in the interactome). The groups were ran separately using functions from the clusterProfiler package^92^ (v4.16.0) with the mouse database org.Mm.eg.db (v3.21.0) using the default thresholds for compareCluster and enrichGO (adjusted p value = 0.05, Benjamini-Hochberg for multiple testing corrections). For comparisons made within PL, any protein bound by at least one hTau protein in PL was included in the background gene set and used in the analysis. The same is true for the BLA comparisons. These two background gene sets were combined for the analyses that compared PL and BLA samples together. The ‘simplify’ function was used to decrease term redundancy for visualization purposes. GO terms not applicable to brain tissue were excluded and some of the remaining top GO terms included in the graphs were shortened or abbreviated for visual purposes. The full, unfiltered ORA output as well as final filtered GO terms are included as supplementary tables. The packages used to visualize the enrichment results included enrichplot (v1.28.4 – treeplots, cnet plots, dot plot), ggtree (v3.16.3, treeplot), and ggplot (dot plots). Identification of mitochondrial-associated proteins in the dataset was carried out using MitoCarta3.0^93^. The Venn diagram was produced using ggVennDiagram (v1.5.7) and the stacked bargraph with ggplot2.

For the phosphoproteomic analysis, all filtered peptides (false discovery rate of < 0.01) were normalized to total peptide abundance over the injected sample. The same strategy described above for determining significance was used with the exception that log2 fold change requirement was decreased to 0.585, given the scarcity of the sample. Abundance values were log2-transformed and globally scaled before creating the heatmap using ComplexHeatmap (v2.24.1). The lollipop graphs and dotplot was created with ggplot2 (v4.0.3). The functional enrichment analyses were carried out as described above.

#### Seahorse analysis

Analysis of the Seahorse data was performed using GraphPad Prism. The Shapiro-Wilk test was used to assess the assay parameters for normality. Basal respiration and ATP production passed and were analyzed using a one-way ANOVA with cell line as the main factor followed by Tukey post hoc tests with corrections for multiple comparisons. Non-mitochondrial oxygen, maximal respiration, proton leak, spare respiratory capacity, spare respiratory %, and coupling efficiency failed the test of normality. They were analyzed using the Brown-Forsythe and Welch ANOVA test followed by Dunnett’s T3 post hoc test with corrections for multiple comparisons.

#### RNA-sequencing analysis

Bioinformatic analysis followed an established pipeline: briefly, the quality of the fastq files was assessed with FastQC v0.12.1 and filtered using fastp v0.24.0 (poly-x tail trimming, poly-X tail trimming, 3’ quality-based tail trimming, Q score >= 15, and length >= 50 bp). Reads were aligned to the latest reference human genome using STAR aligner v2.7.11 with non-canonical splice junction removal and output of unmapped reads and sorted with samtools v1.22.1. UMIcollapse was used to remove PCR and optical duplicates, and alignment quality metrics were assessed using RSeQC v5.0.4 and Qualimap v2.3. MultiQCv1.32 provided an output of the results^94^. Gene expression quantified was performed using featureCounts (subread package v2.1.1; feature identifiers included strand-specific counting, multi-mapping read fractional assignment, exons and three prime UTR, grouped by gene_id). The reference GTF file was used to annotate final gene counts. Graphs were generated in R using the data from the pipeline outlined above. Heatmap and PCA plots were generated using log2-transformed normalized counts (trimmed mean of M-values). Low expression genes, those without a normalized count above 1 in at least 3 samples, were removed. The top 500 most variables genes were selected for hierarchical clustering, and the data was scaled before generating the heatmap. A Gene Ontology over-representation analysis was performed with the genes from each cluster using enrichGO and simplify (cutoff = 0.4). Enriched biological processes were selected for each cluster and added to the heatmap. Gene set enrichment analysis was performed with packages from clusterProfiler including gseGO and simplify (cutoff = 0.5). To look for enrichment in pathways associated with mitochondrial genes, the GSEA function was used with the MitoCarta3.0 database. GO terms not applicable to cells grown in vivo were excluded from the bar graphs and some of the remaining top GO terms are depicted with shortened, condensed versions of the term. The excluded GO terms as well as the original GO term names and their condensed version are included in the supplementary data.

