## Supplementary Figures for "Conserved synaptic modifications by humanized tau through diverse isoform- and mutation-specific intracellular interactions"

**Supplementary Figure 1**

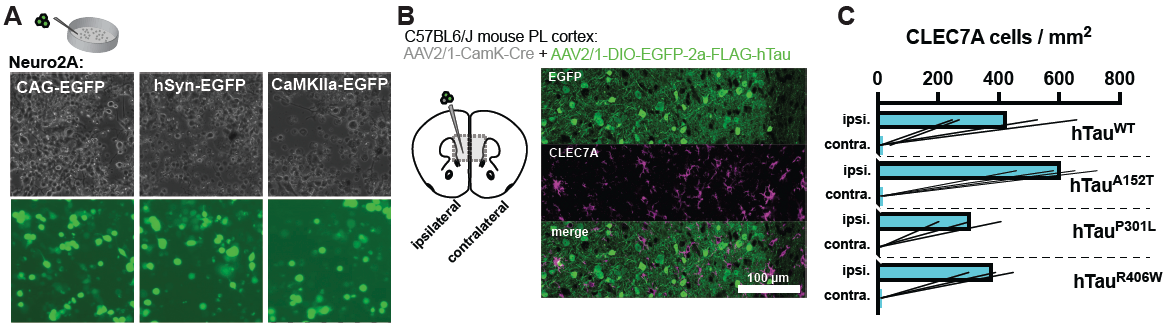

**Supplemental Figure 1. Promoter selection and presence of CLEC7A cells after hTau expression**. **A)** Neuro2a cells were used to transfect different promoters and examine reporter expression. **B)** Co-injection of 0N4R constructs with CaMKII-Cre into mouse PL cortex drove robust EGFP-hTau expression and drove the recruitment of cells expressing the disease-associated microglia marker CLEC7A. **C)** Quantification of cells expressing CLEC7A on the injected (ipsilateral) and noninjected (contralateral) side in PL cortex.

**Supplementary Figure 2**

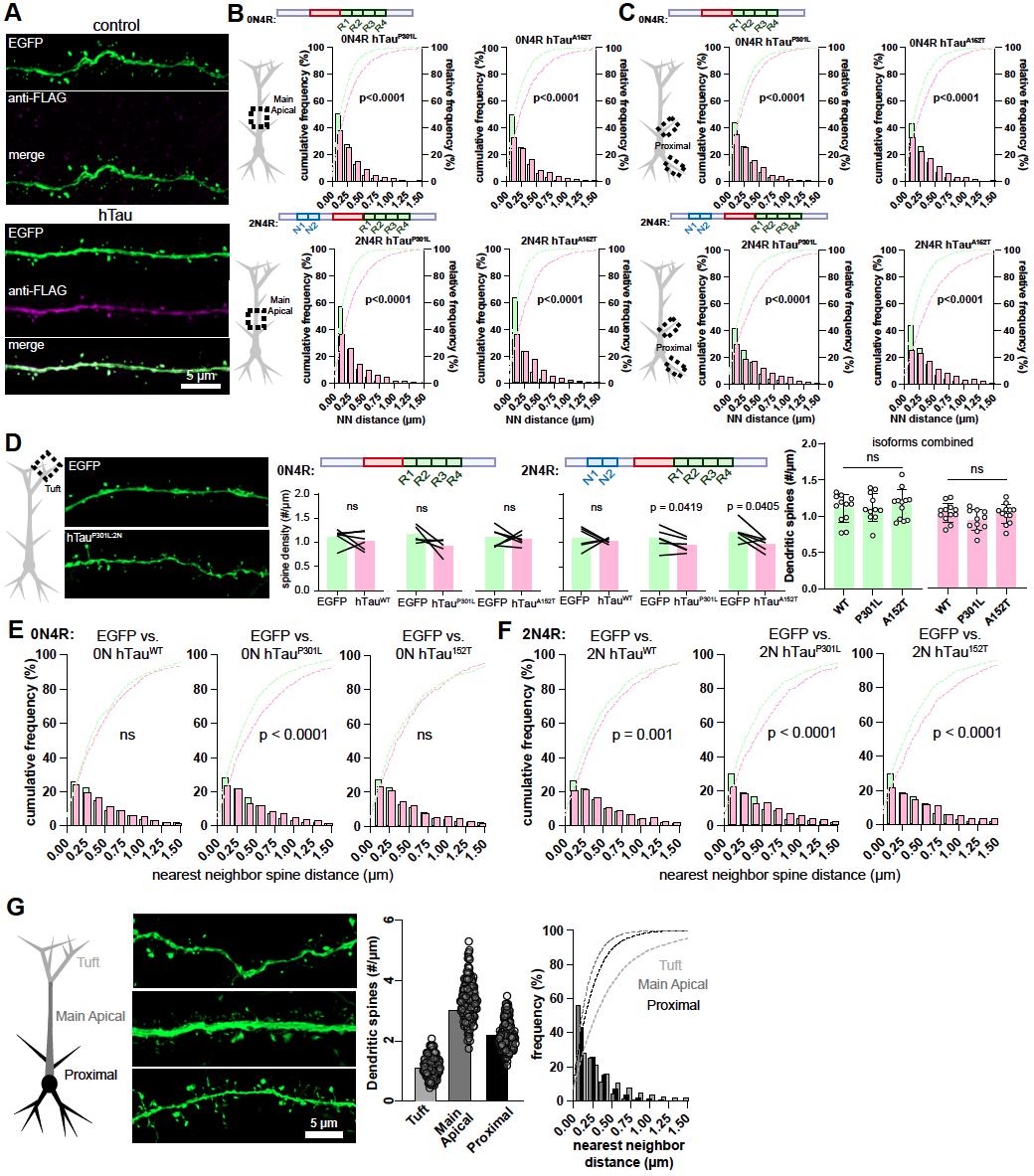

**Supplemental Figure 2. Reduced impact of hTau on synaptic connectivity of distal tuft branches**. **A)** Example of an EGFP-expressing (control) branch segment (top) and a FLAG: hTau-expressing segment (bottom). **B)** Nearest neighbor analyses from the main apical branch for the 0N4R (top) and 2N4R (bottom). **C)** Nearest neighbor analyses from the proximal branches for the 0N4R (top) and 2N4R (bottom). **D)** Within mouse comparisons of distal tuft branches, separated by 0N4R isoforms (left) and 2N4R isoforms (center), and combined across-animal comparisons (right). **E)** Nearest neighbor analyses on tuft branches from 0N4R branches. **F)** Nearest neighbor analyses on tuft branches from 2N4R branches. **G)** Sample branches, branch densities, and nearest neighbor data across EGFP segments from the different domains of PL neurons.

**Supplementary Figure 3**

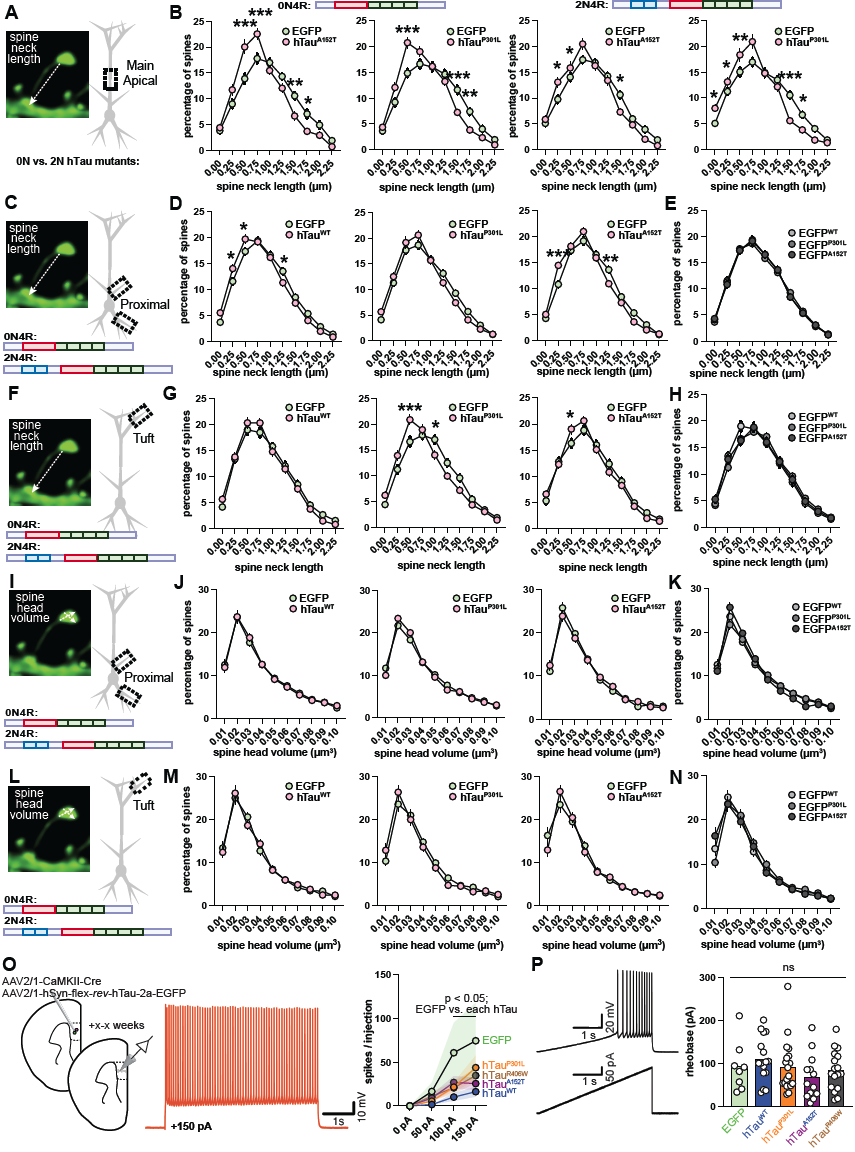

**Supplemental Figure 3. hTau mutant expression alters spine morphology and spike frequency**. **A)** Spine neck lengths were measured from the center of the head to the closest node of the dendritic cable. **B)** Spine neck lengths were taken from segments along the main apical branches from EGFP (control) or EGFP-2a-hTau constructs (30 branches from 0N4R isoforms shown on the left, and 30 branches from the 2N4R isoform shown on the right) from PL-to-BLA neurons. **C)** Spine necks were measures across both isoforms along the proximal branches. **D)** Neck length distributions from hTauWT, hTauP301L, and hTauA152T along proximal branches. **E)** Spine neck lengths from proximal apical oblique and basal branches from the EGFP (control) segments. **F-H)** Same as (C-E) but for spine neck lengths along tuft branches. **I-K)** Same as (C-E) but for spine head volumes along proximal branches. **L-N)** Same as (I-K) but for tuft branches. **O**) Somatic patch-clamp whole-cell recordings were made from excitatory PL neurons expressing EGFP (control) or the 0N4R isoform of hTau^WT^, hTau^P301L^, hTau^A152T^, or hTau^R406W^ (left); example voltage trace of spikes with 150 pA injection (center); and a plot of spike number from increasing current injections. **P)** An example voltage trace in response to a current ramp stimulus (left); estimated rheobase from ramp stimuli from cells expressing EGFP (control) or each 0N4R hTau construct (right).

**Supplementary Figure 4**

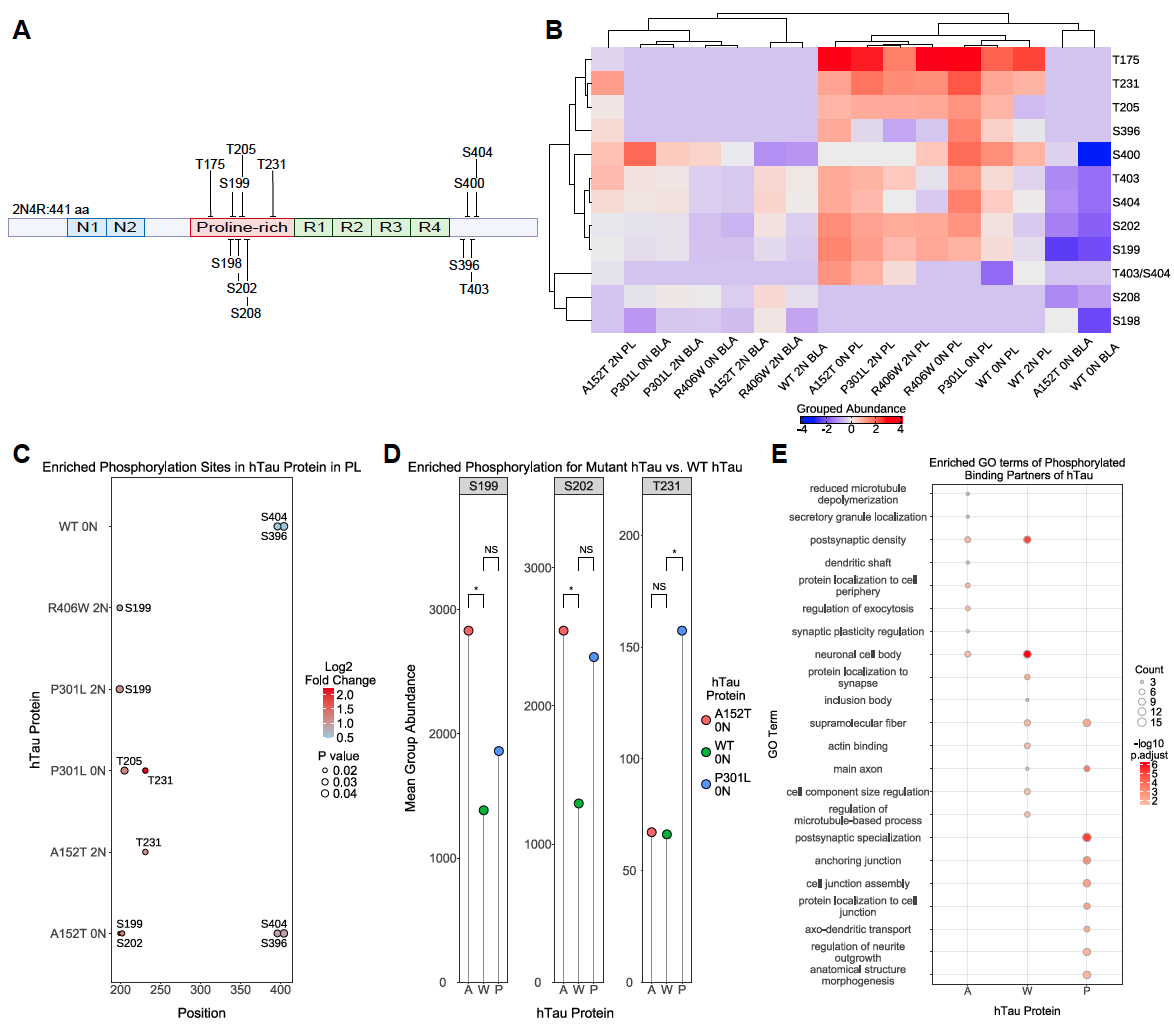

**Supplemental Figure 4. Disease-associated phosphorylation sites identified in hTau samples**. **A)** Schematic of the 2N4R hTau protein with identified phosphorylated residues highlighted. **B)** Hierarchical clustering of the samples based on phosphorylated residue abundance. Color scale indicates the abundance level from high (red) to low (blue). **C)** Graph depicting the enriched phosphorylated sites for the different hTau proteins (y axis) along the length of the hTau protein (x axis). The sites are significantly enriched for compared to the control samples (log2 fold change >= 0.585 and p-value < 0.05). **D)** Lollipop graph indicates three phosphorylated sites (S199, S202, T231) were significantly enriched for in mutant 0N4R hTau samples compared to 0N4R hTau^WT^ (|log2 fold change| >= 0.585 and p-value < 0.05). **E)** Dot plot indicates the top enriched GO terms for the phosphorylated proteins bound to either 0N4R hTau^A152T^, 0N4R hTau^WT^ or 0N4R hTau^P301L^.

**Supplementary Figure 5**

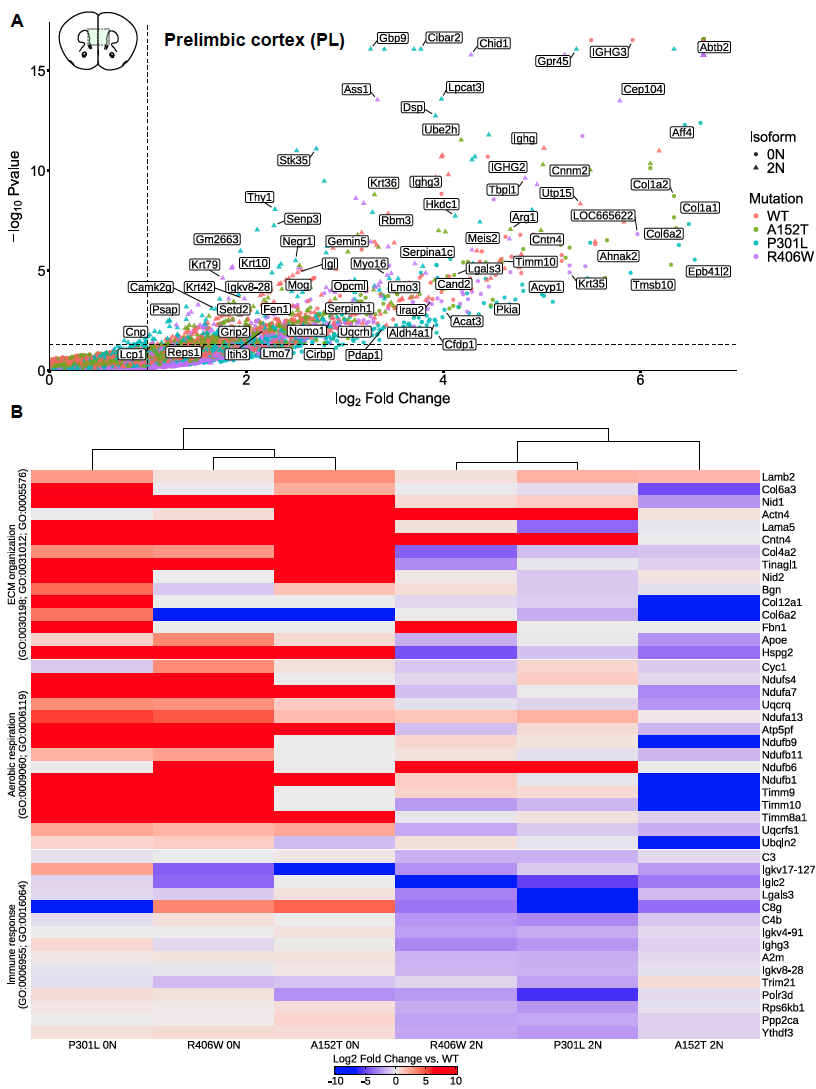

**Supplemental Figure 5. Binding partners of hTau in PL are involved in the extracellular matrix, cellular respiration, and immune response. A)** Volcano plot depicting the significant binding partners of 0N4R hTau proteins (circles) or 2N4R hTau proteins (triangles). The mutations are noted by the different colors. Proteins above the dotted lines were deemed significantly enriched for compared to the control (log2 fold change > 1, adj. p-value < 0.05). **B)** Hierarchical clustering of the samples based on select protein abundances compared to WT. The proteins were enriched for in the depicted GO terms. Color scale indicates the level of relative protein abundance from enriched (red) to depleted (blue).

**Supplementary Figure 6**

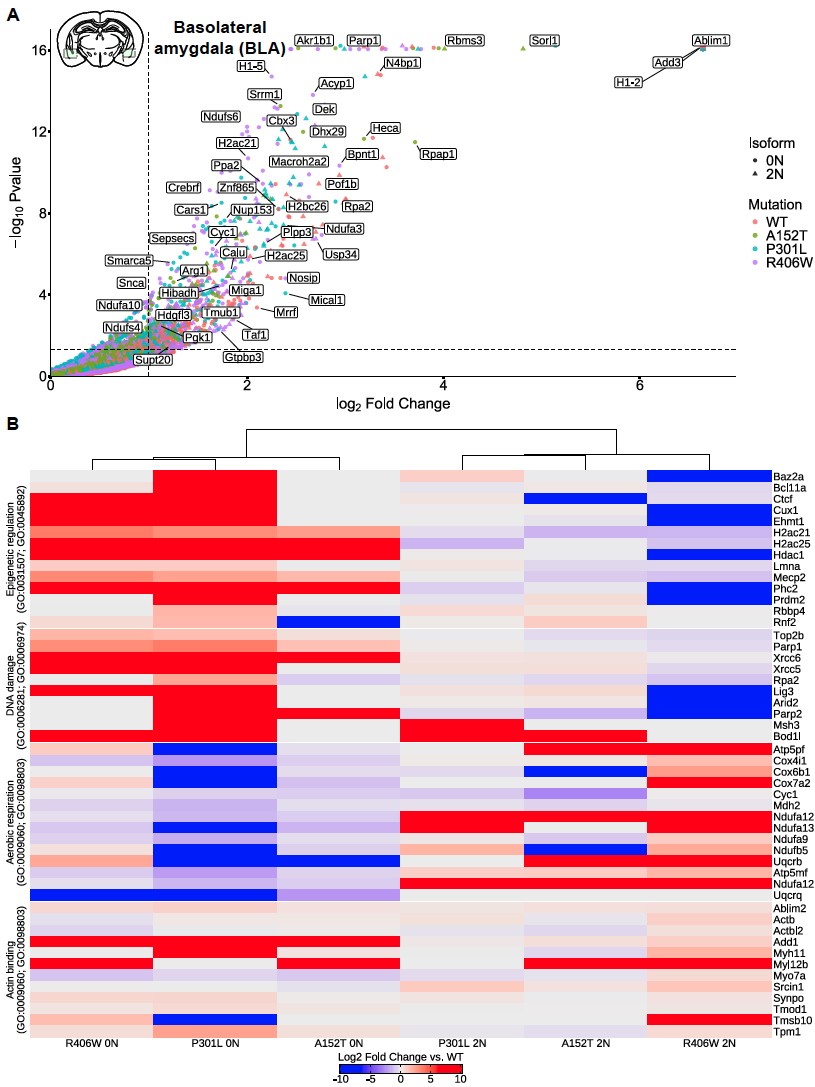

**Supplemental Figure 6. Binding partners of hTau in BLA largely localize to the nucleus and mitochondria. A)** Volcano plot depicting the significant binding partners of 0N4R hTau proteins (circles) or 2N4R hTau proteins (triangles). The mutations are noted by the different colors. Proteins above the dotted lines were deemed significantly enriched for compared to the control (log2 fold change > 1, adj. p-value < 0.05). **B)** Hierarchical clustering of the samples based on select protein abundances compared to WT. The proteins were enriched for in the depicted GO terms. Color scale indicates the level of relative protein abundance from enriched (red) to depleted (blue).

**Supplementary Figure 7**

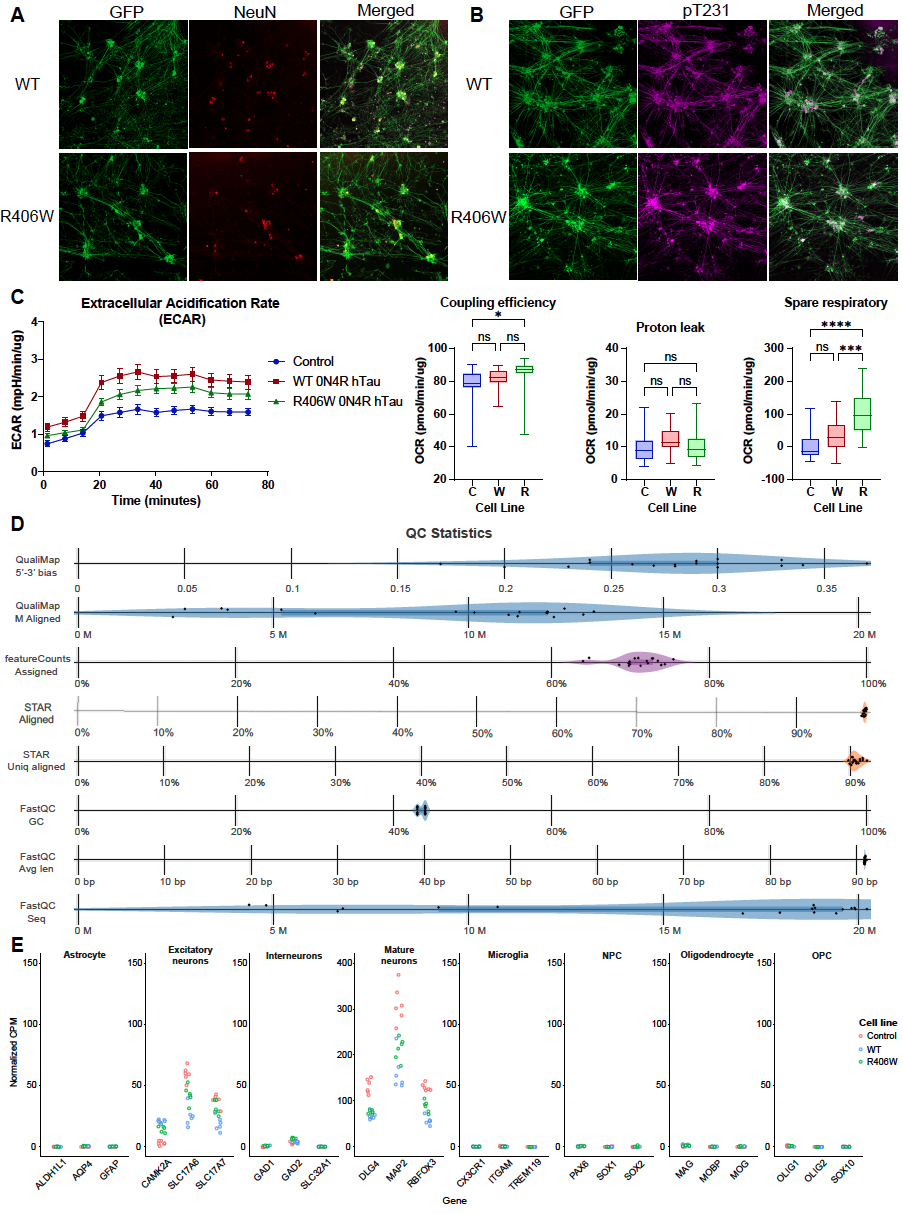

**Supplemental Figure 7. Characterization of the hTau-expressing iNeurons. A-B)** D14 WT and R406W iNeurons expressing EGFP (green) were stained with antibodies (shown in red) for NeuN (A) or pT231 (B). **C)** Additional metrics that were measured for D28 iNeurons during the Seahorse assay including extracellular acidification rate (ECAR), coupling efficiency, proton leak, and spare respiratory capacity. **D)** QC statistics generated with the RNA-sequencing data for the hTau cell lines using the MultiQC tool. The report was provided by Plasmidsaurus. **E)** Strip plot depicting the normalized gene counts (CPM) for the three iNeuron cell lines for marker genes of different cell types.

**Supplementary Figure 8**

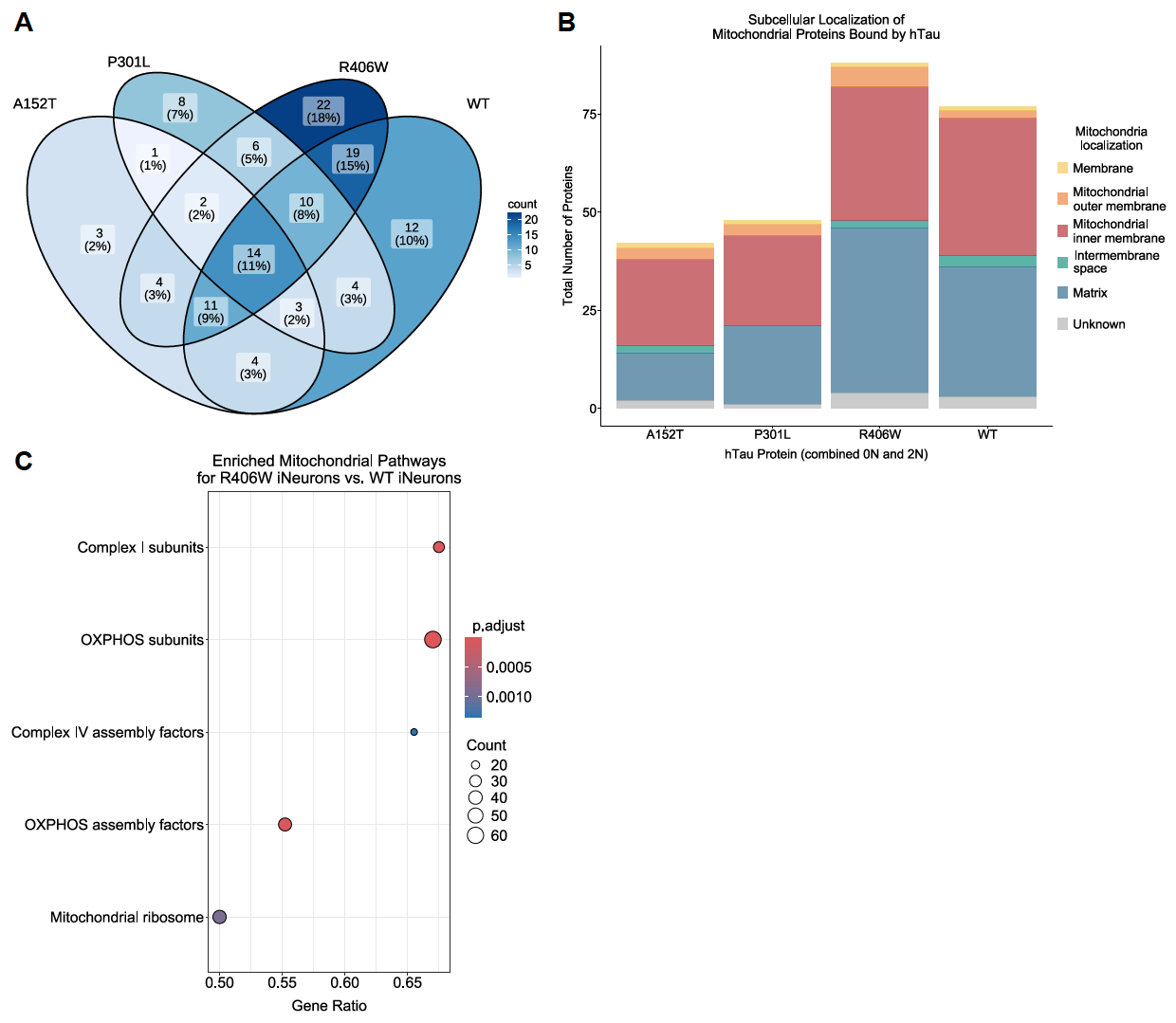

**Supplemental Figure 8. Mitochondrial proteins bound by hTau. A)** Venn diagram illustrating the overlap of mitochondrial proteins in the interactomes of the different hTau proteins. Proteins were classified as being associated with the mitochondria using the MitoCarta3.0 database. The interactomes of the 0N4R and 2N4R hTau proteins are combined for each mutation. **B)** Stacked bar graph demonstrates the subcellular localization of the mitochondrial proteins bound by the different hTau proteins. **C)** Dot plot depicting the five significantly upregulated mitochondrial pathways (GSEA; p value < 0.05) in the R406W iNeurons compared to WT iNeurons. The size of the dots represents the number of genes associated with the pathway and the color scale indicates the adjusted p-value.
